# Ribosome stalling position, spacing, and A-site occupancy impact translation and co-translational mRNA decay in plants

**DOI:** 10.64898/2026.06.12.731700

**Authors:** Sjors van der Horst, Joseph L. Gage, Julia Bailey-Serres

## Abstract

Rate limitation in translational elongation causes ribosomes to pause as they decode an open reading frame (ORF). The triggers and outcomes of these events are poorly understood in plants. Here, we size, map, and quantify footprints of individual (monosome) and closely spaced pairs (disomes) of ribosomes at single-codon resolution to capture stalling and collision events along mRNAs of Arabidopsis and maize. Ribosome footprinting was combined with 5’P-degradome-seq to examine the coincidence of pause events with co-translational decay. Our broad footprint-size selection resolves two monosome conformations and three disome configurations, including monosomes with a vacant or occupied A-site and disomes that are collided or separated by one or two codons. We find positional pausing prevalent at start, stop, and sequential Proline codons, and associated with co-translational processing. These di-Proline pauses are not associated with 5’P peaks. By contrast, brief hypoxia promotes tight stalling of A-site vacant ribosomes at Aspartate codons. This coincides with 5’P peaks on the 5’ side of the stalled ribosome, indicating that rate-limiting decoding can trigger co-translational mRNA decay. Notably, actively transcribed and translated hypoxia-response mRNAs accumulate 1- to 2-codon-separated disomes and are continuously degraded. Comparative footprinting of Arabidopsis and maize reveals that ribosome conformations and codon-specific pausing can be broadly conserved or lineage-specific, as exemplified by pausing on di-Prolines and Conserved Peptide upstream ORFs that regulate production of regulatory proteins. In sum, ribosome stalling at specific codons coupled with ribosome A-site occupancy and disome spacing modulates protein production and co-translational mRNA decay in plants.

**Significance Statement:** The translation of mRNAs into protein relies on tRNAs to deliver amino acids to ribosomes that incorporate these building blocks into the growing polypeptide chain. By monitoring ribosomes as they progress from one codon to the next, we identified ribosome “traffic jams” with different characteristics, causes, and consequences. We find that decoding of certain codons and codon pairs guides polypeptide synthesis and mRNA recycling. The environment can influence this regulation. A comparison of plant species separated by nearly 150 million years highlights deep conservation in positional ribosome stalling patterns that limit translation of mRNAs encoding a suite of regulatory proteins.

## Introduction

The translation of mRNA into protein by ribosomes is an ancient and essential cellular mechanism. In plants, it is one of the most energy consuming processes and is therefore a limiting factor in cell growth (1, 2). mRNAs are decoded by multiple ribosomes simultaneously in polysome complexes, enabling high levels of protein synthesis. The initiation phase of translation is selective and rate limiting, but elongation can become discontinuous if a ribosome pauses, causing a ribosome collision by the trailing (more upstream) ribosome (3). The causes and effects of ribosome stalling in eukaryotes are varied. Despite demonstration in plants of translational regulation in the context of developmental programs and environmental cues (4, 5), there has been limited assessment of the physical distinctions and mechanistic impacts of ribosome stalling events.

In eukaryotes, positional ribosome stalling guides a number of co-translational processes that influence protein production. These can include nascent polypeptide folding, processing, ER secretion, mitochondrial import, and protein complex assembly (3, 6). Upstream open reading frames (uORFs) typically limit the frequency of initiation of the downstream main (m)ORF, which often encodes a regulatory protein. A range of hormone, nutrient, abiotic and biotic stress conditions modulate uORF-guided stalling in plants (5, 7). A number of uORFs that encode conserved peptide (CP) sequence, are known to determine conditional ribosome pausing mediated by levels of specific metabolites (8, 9). A well characterized example is the sucrose-mediated stalling on a CPuORF that precedes the mORF of genes that encode S1-group *BASIC LEUCINE ZIPPER11* (*bZIP11*) transcription factors of Arabidopsis (10). Cryo-electron microscopy has shown that stalling of ribosomes near the CPuORF stop codon involves a physical interaction between the nascent peptide and sucrose within the polypeptide exit tunnel of the ribosome (11). In seedlings grown under standard conditions, ribosomes accumulate just 5’ of the CPuORF stop codon on S1-*bZIP* mRNAs (12). Exposure to brief hypoxia, a low energy stress, decreases the number of ribosomes stalled on these uORFs, concomitant with an increase in the proportion of ribosomes on the mORF. The stalling and structural characteristics of ribosomes translating CPuORFs has not been broadly examined.

Ribosome stalling can contribute to mRNA turnover. In mammals and yeast, stalling that results in collision between ribosomes can activate a no-go decay (NGD) process that triggers endonucleolytic cleavage of the transcript, followed by degradation in the 5’ to 3’ direction by a conserved exonuclease (XRN) (13) and in the 3’ to 5’ direction by the SUPERKILLER complex (13–18). In Arabidopsis, the 5’ to 3’ exonuclease XRN4 degrades 5’ decapped mRNAs, producing an RNA with a free 5’ monophosphate (5’P) terminus (19). The NGD pathway is integrated into the Ribosome-Associated Quality Control (RaQC or RQC) pathway that clears the nascent polypeptide and recycles the ribosome subunits. This is initiated by ubiquitylation of two small ribosomal subunit (40S) proteins of the stalled ribosome, followed by dissociation of the 40S and 60S ribosome subunits, facilitating ubiquitin-mediated turnover of the nascent polypeptide (3). Orthologs of RaQC proteins have been recognized in Arabidopsis and other plants (20, 21). Notably, not all ribosome stalling events determine the same outcome. In mammals, cryo-EM imaging has shown that persistent stalling accompanied by physical ribosome collisions can promote co-translational mRNA decay or boost ribosome elongation past pause sites, a phenomenon dubbed ribosome cooperativity (22). Finally, positional ribosome stalling is essential for the unconventional cytoplasmic removal of conserved introns in animals and plants (23, 24).

In plants, mRNA decapping followed by XRN4-mediated 5’ to 3’ decay modulates mRNA abundance in diverse developmental and environmental contexts, ranging from acute heat and circadian regulation to seedling development (25–27). Decay intermediates with the tell tale 5’P termini are monitored using modified RNA-seq methods. So called degradome-seq methods also are used to identify 5’ cleavage sites associated with silencing by microRNAs and small RNAs. Several degradome studies on plants report a 3-nt codon-based periodicity in 5’P reads that is consistent with XRN degrading mRNAs associated with translating ribosomes (28, 29, 30). This is supported by the loss of 3-nt periodicity in the RNA degradome of an enzymatically active XRN4 mutant that lacks a domain necessary for association with ribosomes (31).

To gain insight into ribosome pausing and collision events in plants, we performed single and paired (disome) ribosome footprint isolation and sequencing (monosome-seq and disome-seq) on Arabidopsis\ seedlings grown under standard conditions and following brief hypoxia. This was complemented with traditional RNA-seq and 5’P-degradome-seq to monitor mRNA abundance and co-translational decay, respectively. We resolve two monosome conformations and three disome spacing configurations: ribosomes with or without an aminoacyl (aa)-tRNA bound to the ribosome A-site and disomes that are tightly packed or separated by one or two codons. The data recognize position-, sequence- and condition-specific stalling associated with certain codons and codon pairs. Ribosome pausing can position polypeptides for processing and tightly packed disomes can trigger co-translational decay. Finally, we show that ribosome conformation, spacing and pausing behavior that modulate production of proteins are conserved on orthologous uORFs and mORFs of Arabidopsis and maize, with notable distinctions.

## Results and Discussion

### High-resolution footprinting resolves aa-tRNA vacant and occupied A-sites of ribosomes and spacing variation between closely packed ribosomes

To explore ribosome stalling in plants, we performed monosome-seq and disome-seq on flash-frozen Arabidopsis ecotype Col-0 seedlings grown in air or treated with hypoxia for 2 h. Disome-seq is similar to ribo-seq (referred to here as monosome-seq), where, after treating polysomes with RNase I, transcript fragments protected from degradation by ribosomes (ribosome footprints) are extracted and processed for short-read sequencing (**Fig. 1*A***). Monosome-seq protocols use high concentrations of RNase I to achieve digestion of mRNA to the edges of the 80S ribosome. Although this improves the accuracy in mapping the most 5’ nucleotide of the ribosome footprint relative to the ribosome aa-tRNA P-site, it can convert closely spaced disomes into monosomes (6, 32), masking ribosome collisions and thereby reducing the ability to detect translational pausing and stalling events. We optimized RNase I digestion conditions to avoid over digestion of disomes, preserving both monosome and disome peaks (***SI Appendix,* Fig. S1**).

**Fig. 1.**
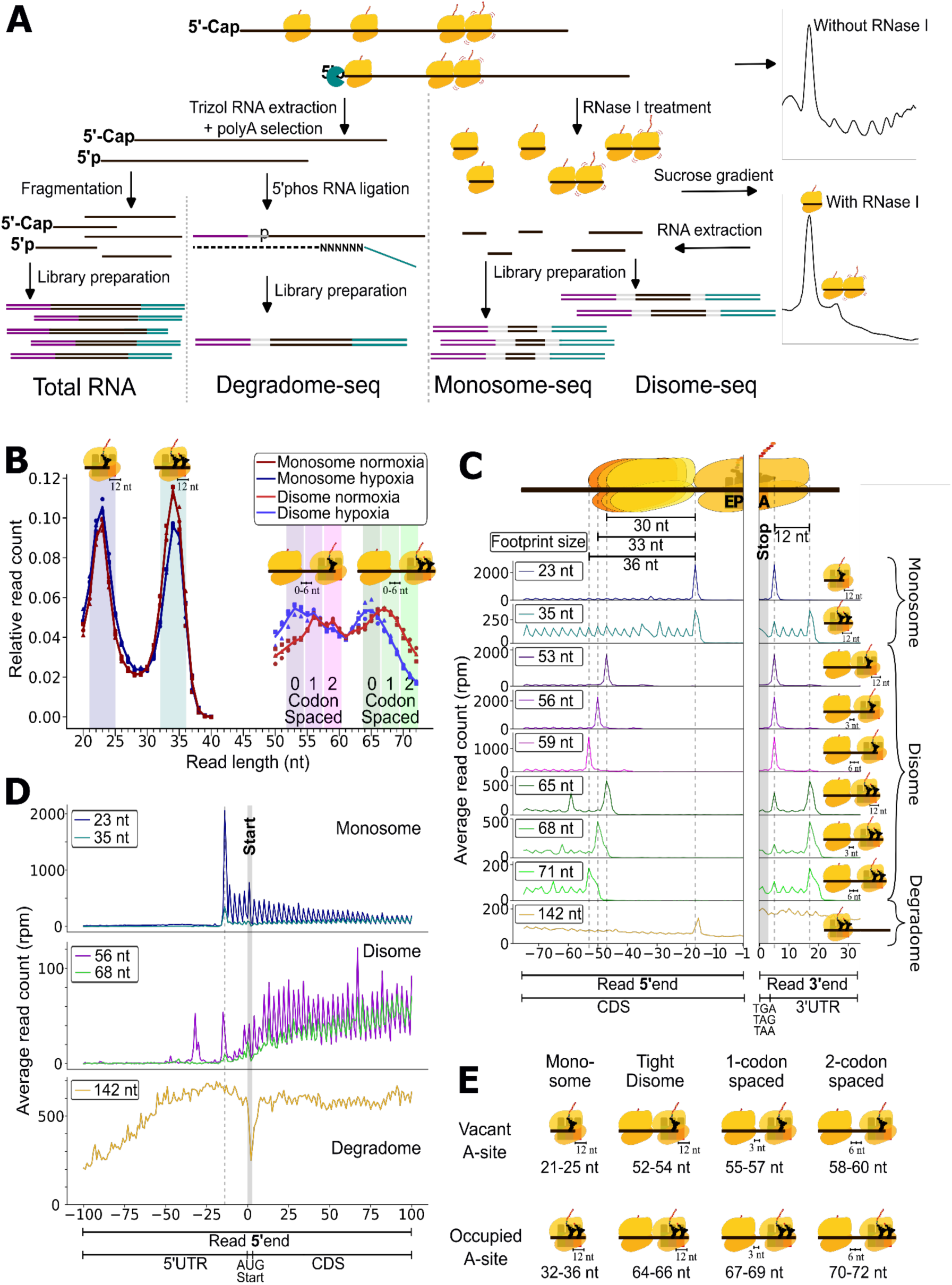
Broad examination of mono- and di-ribosome footprints exposes distinct ribosome conformations and disome packing density. (***A***) ‘Total RNA’ and ‘Degradome-seq’ (5’P) library preparation involved isolating polyA RNA from 7-d old Arabidopsis seedlings by use of oligo(dT) affinity purification followed by fragmentation or 5’ phosphate linker ligation, respectively. ‘Monosome-seq’ and ‘Disome-seq’ libraries involved cell lysis, RNase I treatment, and monosome and disome separation by use of sucrose density gradients, followed by gel fractionation. See Material and Methods for details. (***B***) Average read length distribution of monosome-seq and disome-seq footprints of seedlings treated with or without 2 h of hypoxia. (***C***) Read 5’ and 3’ end distribution upstream or downstream of the stop codon, respectively, for different library types and read lengths. Schematic of ribosomes at the top and on the right represent the respective positions when the A-site of the leading ribosome is at a stop codon. Reads per million (rpm) averages of 3 bioreplicates are shown. (***D***) Read 5’-end distribution around the start codon. (***C,D***) *x*-axis units are distance from indicated codon in nt. (***E***) Overview of two distinct single ribosome conformations (A-site vacant [smaller footprint]; A-site aa-tRNA occupied [larger footprint]) and six configurations of disomes separated by 0 to 2 codons (0 to 6 nt). Ribosome tRNA binding site depiction is from left to right, as shaded rectangles, along the mRNA (5’ to 3’). E, Exit site; P, Peptidyl-tRNA binding site; A, Aminoacyl-tRNA binding site. tRNAs are shown as black triangles in the A or P tRNA binding site, and the tRNA-tethered nascent polypeptide as a red chain.

Typically, monosome-seq is performed on footprints ranging from 27 to 36 nt, corresponding to ribosomes with an aa-tRNA-occupied A-site (33). We broadened the size of fragments collected from 20 to 38 nt in order to capture the smaller footprints of ribosomes lacking an aa-tRNA bound to the A-site. We refer to these as vacant A-site ribosomes. This captured prominent monosome footprint size peaks of approximately 23 and 35 nt, corresponding to vacant and occupied A-site ribosomes, respectively (**Fig. 1*B***). The 5’ ends of the 23 and 35 nt footprints map 17 nt upstream of the 5’ nucleotide of the ribosome P-site, indicating a similar ribosome conformation at the mRNA channel exit site. The 3’ ends of these footprints peak 5 and 17 nts downstream of the first base of the stop codon, respectively (**Fig. 1*C***). This confirms a 12 nt difference in footprint size between aa-tRNA vacant and occupied A-site ribosomes.

Thus, our optimized polysome digestion and protected fragment collection strategy captures two ribosome subpopulations due to differences in protection at the 3’ end of the footprint. This is consistent with the short and longer footprints of vacant or occupied ribosome A-sites described for yeast and mammals (33, 34). The predominance of 23-nt footprints at Arabidopsis stop codons supports the hypothesis that this footprint arises from ribosomes awaiting the binding of Release Factor 1 (eRF1) to the A-site, enabling hydrolysis of the bond between the polypeptide and the final aa-tRNA (**Fig. 1C**). Next, we asked if the same ribosome conformations are observed in closely spaced pairs of ribosomes.

To monitor disomes we isolated, sequenced, and analyzed disome footprints ranging from 50 to 72 nt, resolving peaks of 53, 56, 59 nt and ∼67 nt (**Fig. 1*B***). Using the termination codon region as a guide, we find the 3’ ends of the shorter footprints peak 5 nt downstream of the stop codon, as observed for vacant A-site monosomes, and the longer footprints (65, 68 and 71 nt) peak 17 nt downstream, as observed for the A-site occupied monosomes (**Fig. 1*C***). The 5’ ends of disomes peak at 47, 50 or 53 nt upstream of the stop codon for both the short and long footprints. Notably, the same disome footprint sizes are found within coding regions. From these data we conclude that disomes can be tightly packed (52-54 and 64-66 nt), or separated by one (55-57 and 67-69 nt) or two codons (58-60 and 70-72 nt), with the downstream ribosome having either a vacant or occupied A-site (**Fig. 1*C,E***). We refer to the tightly packed disomes as collided ribosomes and the more loosely packed disomes as paused ribosomes. Ribosomes could crowd because of high rates of initiation or when awaiting a rate-limiting event (*e.g*., aa-tRNA or eRF1 binding, peptidyl transferase or translocation reactions, co-translational protein folding or processing).

We reasoned that tightly packed disomes resulting from prolonged pausing or collision could trigger co-translational decay of mRNA. To investigate this, the same samples were processed for 5’P RNA degradome-seq. This method maps 5’ termini with a free monophosphate group generated by 5’ decapping and 5’ to 3’ digestion by XRN4 or through endonucleolytic cleavage, such as that mediated by miRNA or sRNA silencing. Degradome-seq reportedly captures the most 5’ position protected by a ribosome that is “followed” by XRN4 in Arabidopsis (26, 29, 31) and Xrn1 can be positioned at the mRNA exit channel of the last (most 5’) translating ribosome of an mRNA in yeast and mammals (3). Our data resolve very clear 3-nt codon periodicity in 5’P termini on ORFs and a peak 16-17 nt upstream of stop codons (**Fig. 1 *C,D*; *SI Appendix,* Fig. S2**). This confirms that 5’ to 3’ co-translational mRNA decay operates along translating mRNAs and can extend to a terminating ribosome.

In sum, our method distinguishes two monosome conformations and six disome configurations providing insight into A-site occupancy as well as the distance between disomes separated up to two codons (**Fig. 1E**). By contrast, combined monosome- and disome-seq studies of yeast and vertebrates have not captured the same suite of footprint sizes resolved here and were not coupled with degradome data. Footprints from ribosomes with vacant A-sites were often absent, likely due to a narrower selection of fragment sizes (35, 36). Although some studies report disomes with one-codon spacing and occupied A-sites, disomes with one-codon spacing and empty A-sites or those with two-codon spacing are not reported (6, 32, 36). A possible explanation for the underrepresentation of less packed disomes in these studies is differences in the extent of RNase digestion. Higher RNase levels or longer reaction times may cleave mRNA between closely spaced ribosomes, and overly aggressive digestion may degrade ribosomes entirely, converting trisome footprints into disomes, and disomes into monosomes, resulting in “stacking” of footprints that are characterized by contiguous, back-to-back read placement (6, 32, 36). Our datasets allow us to compare A-site occupancy and disome spacing patterns across ORFs, and to identify stalling events associated with mRNA decay.

### Translational elongation ramps up

Beginning with the start codon, we observe an enhanced peak of 23 nt vacant A-site footprints, with a 5’-end mapping to −17 nt from the +1 position of the coding sequence. These vacant A-site ribosomes progressively decline to equilibrium about 75 nt (25 codons) downstream, as the 35 nt footprints of A-site occupied ribosomes trend upwards (**Fig. 1*D***). At this point, levels of 23 and 35-nt footprints are similar and accompanied by well matched levels of one-codon separated disomes with or without an A-site tRNA. We reason that the presence of A-site occupied and vacant ribosomes at the start site reflects longer Met-tRNA residence time or slower ribosomal peptidyl transferase activity and translocation as the initiating ribosome transitions to the first round of elongation. Despite the shift from vacant to occupied A-site ribosomes in this region, the 5’P reads indicative of co-translational decay are stable, except within the first few codons where they are nearly absent.

Studies on yeast and animals report that elongation rates increase over the first 50 codons, a phenomenon referred to as the translational ramp. Our resolution of a transition from predominantly vacant to occupied A-site ribosomes in the first 25 codons is consistent with a translational ramp. Causes of a ramp are debated. One proposal is that rare codons in this region limit ribosome collisions (37), another is that early crowding of ribosomes promotes rather than suppresses translational productivity (38, 39). The ramp region could overlap with co-translational protein targeting and processing events, such as engagement of the signal sequence of a nascent polypeptide with the signal recognition particle (SRP) complex and docking on the ER (23, 35). This occurs either before or after the signal sequence emerges from the ribosome (40, 41). Yet, we find no clear evidence of early pausing on mRNAs encoding the ensemble of secreted and organelle-target proteins (***SI Appendix,* Fig. S3**). Other factors that could determine a slow start in translation of individual mRNAs include co-translational N-terminal processing (acetylation, myristoylation, methionine excision, N-degron modifications) or nascent protein association complex engagement (20). It could be informative to include disome-sequencing in the analysis of mutants that affect N-terminal processing and protein targeting to resolve the role of ribosome state dynamics and links to mRNA turnover in these processes.

We were intrigued to find that a small proportion of all mapped monosome and disome reads map to 5’ and 3’UTRs (2.2 ± 0.06% and 1.4 ± 0.10%, respectively) (***SI Appendix,* Fig. S4*A,D***). Ribosomes mapping to 5’UTRs are predominantly those with vacant A-sites (***SI Appendix,* Fig. S4*A***) and may correspond to reads on uORFs found in an estimated 31% of Arabidopsis 5’ leaders (42). The bias towards A-site vacant ribosomes in 5’UTR footprints seems unlikely to reflect the presence of rare codons in uORFs, as these are not preferred in expressed Arabidopsis uORFs (43). Another explanation for the vacant A-site bias on 5’UTRs could be the short length of uORFs overrepresents start site, translational ramp, and stop codon-localized ribosomes, as compared to much longer mORFs. These findings highlight the importance of sampling shorter ribosome footprints when investigating translation events in 5’UTRs. Also the small proportion of Arabidopsis ribosome footprints on 3’UTRs is reminiscent of observations for cultured Drosophila and human cells (44). Ribosome traversal of the 3’UTR can be due to stop codon readthrough, incomplete ribosome dissociation during the recycling phase, or alternative splicing generation of non-stop mRNA.

### Identification of stall sites at single codon resolution

We reasoned that a comparison of air exposed (normoxic) and briefly hypoxic (2 h) seedlings would be appropriate to investigate conditional changes in monosome conformations and disome spacing. In our hands this stress reduces cellular ATP levels by about 50% with a proportionate reduction in global translation of cellular mRNAs, as determined by quantitative polysome profiling (12, 45). The rapid and reversible stress response involves the transient sequestration of a large proportion of mRNAs into biomolecular condensates marked by the stress granule protein OLIGOURIDYLATE BINDING PROTEIN 1C (UBP1C) (46). UBP1C condensation and nucleus retention of mRNA enables the selective translation of newly synthesized core hypoxia-responsive gene transcripts (47). We found that the monosome footprint size distribution of hypoxic seedlings does not differ appreciably from those grown in air. The disome footprint distribution under hypoxia shifts towards tightly stacked disomes (53 and 65 nt), suggesting a conditional increase in ribosome density and collisions (**Fig. 1*B*; *SI Appendix,* Fig. S4*B***).

To explore the relationship of disome formation to cotranslational decay, the decoding of individual amino acid codons was investigated by comparison of ribosome footprint and degradome data. For each codon, a peak score was calculated by dividing the number of reads at the predicted A- or P-site of a codon by the average of the surrounding codons as described in the Methods (**Fig. 2*A***). A peak score >1 indicates an increased number of ribosomes at that A- or P-site, indicating slower decoding of the codon, whereas a score <1 indicates faster decoding. Regardless of condition, the start (initiation) and stop (termination) codons cause the most pronounced stalling based on their high peak scores for the monosome and disome, and degradome-seq data. Stop codon stalling occurs when the codon is in the A-site, whereas start codon stalling occurs when positioned in the P-site. The lack of disomes at the start codon reflects absence of a downstream ribosome, whereas the drop off in degradome reads may reflect our intentional omission of cycloheximide from the degradome extraction buffer, as cycloheximide may artificially increase the start codon peak by blocking elongation but not initiation during tissue lysis (48). We note that methionine (Met/M) codons, other than the annotated initiation codon, show a higher peak score in the monosome data (**Fig. 2*A* and Dataset S1**). These could be alternative initiation sites. The clarity of this single-codon resolution of ribosome confirmation and spacing motivated us to consider if other codons and peptide motifs associated with slow decoding were associated with mRNA decay.

**Fig. 2.**
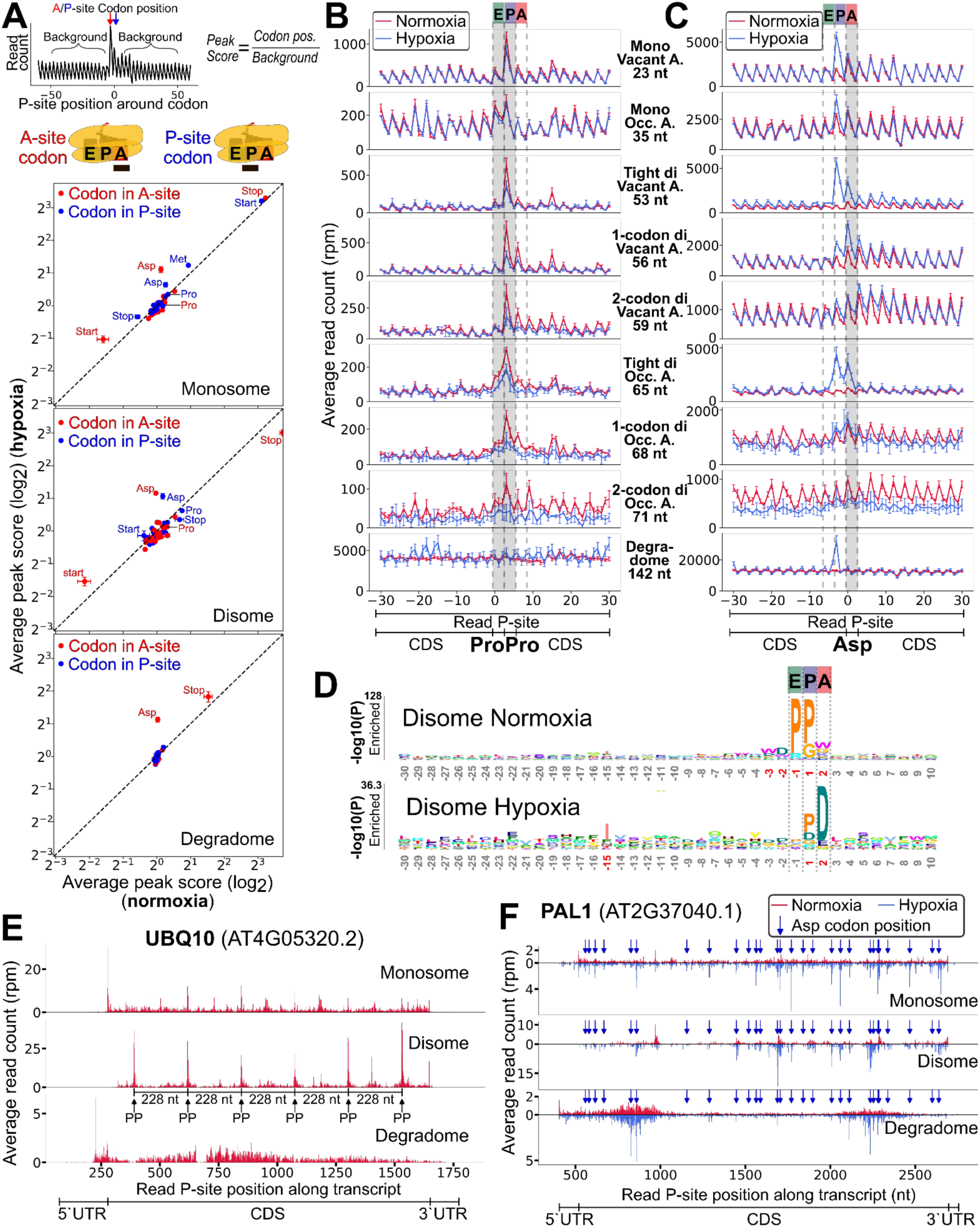
Slow di-Pro codon decoding is associated with ubiquitin monomer production whereas hypoxia-induced slow Asp codon decoding is associated with signatures of co-translational decay. (***A***) Log_2_ of average peak score (read count at codon A-site [red] or P-site [blue] divided by background) of the 20 amino acid codons (sets) and the start and stop codons of hypoxia (*y*-axis) or normoxia (*x*-axis) data (input data: **Dataset S1**). (***B,C***) Predicted P-site monosome, disome or degradome RPM distribution around di-Pro (***B***) and monoAsp (***C***) encoded codons of different RNA-seq library types and read lengths. *x*-axis units are distance from indicated codon in nts. (***D***) Probability logo based on stalling sequences (n= 838, normoxia and n=142 hypoxia genes), positions highlighted in red indicate significantly enriched positions (Bonferroni corrected p-value < 0.01). (***E, F***) P-site position RPM on *POLYUBIQUITIN10* (*UBQ10,* AT4G05320.2) and *PHE AMMONIA LYASE 1* (*PAL1*; At2G37040.1) exemplifying di-Pro stalls (***E***) or hypoxia-induced monoAsp stalls, indicated with blue arrows (***F***). (***A-C***) Error bars represent standard deviation from three biological replicates.

### Polyproline-induced ribosome stalling is observed in monosome- and disome-seq but not in the 5’P degradome-seq data

Previous research shows that consecutive Prolines (Pro/P) are rate limiting in elongation due to the requirement of eukaryotic Initiation Factor 5A (eIF5A) for di-Pro peptide bond formation (49). A peak score >1 is observed for Pro codons in the P-site in both monosome- and disome-seq data, independent of the specific Pro codon used (**Fig. 2*A*; *SI Appendix,* Fig. S5*C-F***). This effect is even stronger for di-Pro codons, especially when situated in the ribosome E- and P-sites (***SI Appendix,* Fig. S6**). A deeper analysis reveals that di-Pro E+P-site stalling is observed under both normoxia and hypoxia in both ribosomal conformations, although the vacant A-site (tight) configuration predominates (**Fig. 2*B*, 2*D***). There is no evident increase in 5’P degradome peak score at di-Pro pause sites (**Fig. 2*B***), revealing that these events are not typically associated with mRNA decay.

High ribosome density was previously on mRNAs encoding Pro-rich proteins (12, 50). A transcript with multiple identified di-Pro stall sites is *POLYUBIQUITIN 10* (*UBQ10*). *UBQ10* encodes six monoubiquitin repeats that are processed into monoubiquitin units by deubiquitinases (DUBs) (**Fig. 2*E***). The stall sites on *UBQ10* mRNA are separated by 228 nt (76 codons), which exactly span a single UBQ monomer. Disome stalling occurs at the conserved sequence PPDQQ, with di-Pro codons in the E-, P-, and an Asp/D codon in the A-site of the ribosome. The same period and pattern of stalling is observed on *UBQ3* and *UBQ14* mRNAs (***SI Appendix,* Fig. S7*B-C***), and on polyubiquitin mRNA in yeast (36). This indicates conserved di-Pro-mediated pausing during translation of *UBQ* mRNA guides or reports DUB activity (***SI Appendix,* Fig. S8*A***).

*UBQ* mRNA and other di-Pro-related pauses do not correlate 5’P degradome peaks despite clear ribosome position tracking, based on 5’P site periodicity and degradome peaks near the stop codon under both normoxia and hypoxia conditions (**Fig. 2*A***). Notably, the Asp/D codon in vacant A-site of the stall motif PPDQQ of *UBQ10* does not enhance stalling under hypoxia, indicating that the di-Pro codons in the P and E sites are the major factor causing these stalls (***SI Appendix,* Fig. S5*A-B*; S7*A***). Thus, di-Pro can serve as a conserved positional translational pause site to facilitate co-translational polypeptide conformational changes or processing events that are not associated with 5’ to 3’ mRNA decay.

### Decoding of aspartate codons under hypoxia

We find that hypoxia monosome and disome increases ribosome occupancy at Asp codons concomitant with elevated 5’P termini of the stalled ribosome (−13 from the P-site) (**Fig. 2*A***). This effect is independent of the specific Asp codon (***SI Appendix,* Fig. S5*A-B***) and primarily when the Asp codon is in an A-site vacant ribosome in monosomes (25 nt) and tightly stacked (collided, 53 nt) disomes (**Fig. 2*C***). This slow Asp decoding appears to be conditionally rate limiting, promoting stalling and collision with upstream ribosomes. *PHE AMMONIA LYASE 1* (*PAL1*) and *DROUGHT-REPRESSED 4/KUNITZ TRYPSIN PROTEASE INHIBITOR 11* (*DR4/KT11*) illustrate hypoxia-induced Asp codon stalling (**Fig. 2*F*; *SI Appendix,* Fig. S7*E***).

Hypoxia enhanced Asp codon stalling correlates with significant reductions in Asp levels in roots and shoots of stressed Arabidopsis seedlings grown and treated in the same set-up (51). The decline in Asp under hypoxia involves its consumption to augment generation of NAD^+^ via a modified TCA cycle coupled with substrate-level ATP production (52, 53). The elevation of tight disomes at vacant A-site Asp codons could coincide with a decline in the Asp-tRNA pool (***SI Appendix, Fig S8B***). Unlike di-Pro stalling, monoAsp stalling can coincide with 5’P transcript termini, as seen conditionally on *PAL1* and *DR4* mRNA during hypoxia (**Fig. 2*F*; *SI Appendix,* Fig. S7*E***).

As an alternative approach to identifying determinants of ribosome stalling, pronounced monosome and disome footprints indicative of stall sites were identified on individual mRNAs and the predicted P-site codon sequences and their surrounding codons were examined. Stall sites were identified as peak scores of at least 20 and supported by ≥5 reads in each of three replicates for both normoxia (n=838) and hypoxia (n=142) (***SI Appendix,* Fig. S9*A* and File S1**). A probability plot of amino acid sequences reconfirms significant enrichment of di-Pro motifs under normoxia and monoAsp codons under hypoxia (**Fig. 2*D*; *SI Appendix,* Fig. S9*A***). Although we find no association between 5’P read peaks (n=1157) and specific codons in the E, P or A site of ribosomes under normoxia, vacant ribosomes awaiting Asp tRNA binding stand out at monosome, disome and degradome peaks (n=1610) under hypoxia.

We hypothesize that conditional reduction in Asp-tRNA levels promote vacant A-site ribosomes to stall at monoAsp codons, fostering collision by upstream ribosomes and activation of 5’ to 3’ co-translational decay (***SI Appendix,* Fig. S8*B***). Disome footprinting with mutants deficient in NGD or transgenes that replace Asp collision sites with Glutamate/Glu codons could be used to test this hypothesis. It is interesting to consider that positional Asp codon stalling may contribute to mRNA decay within the hypoxic microenvironment of apical shoot meristems of Arabidopsis (54).

### Global coordinated changes in transcriptome, monosome, disome and degradome upon hypoxia

To investigate the extent of coordinated versus distinct gene expression regulation induced by hypoxia, we clustered differentially regulated genes (DRGs; |log_2_ Fold Change| > 1; p adj < 0.01) in at least one of the four sequencing dataset types into clusters and analyzed gene ontology (GO) enrichment (**Fig. 3*A*; *SI Appendix,* Fig. S10**). Two-thirds of the core hypoxia response genes (HRGs) (47, 55) are upregulated at the polyA mRNA, degradome, monosome, and disome level (clusters 1-3) (**Fig. 3*A,B*; *SI Appendix,* Fig. S11*A-B* and SI Dataset 2A)**. The high degradome signal demonstrates these transcripts are high-flux, with coordinated synthesis and decay during hypoxia. Genes in clusters 1-4 are broadly associated with environmental interactions including response to hypoxia, chitin, abscisic acid, alcohol, and salicylic acid.

**Fig. 3:**
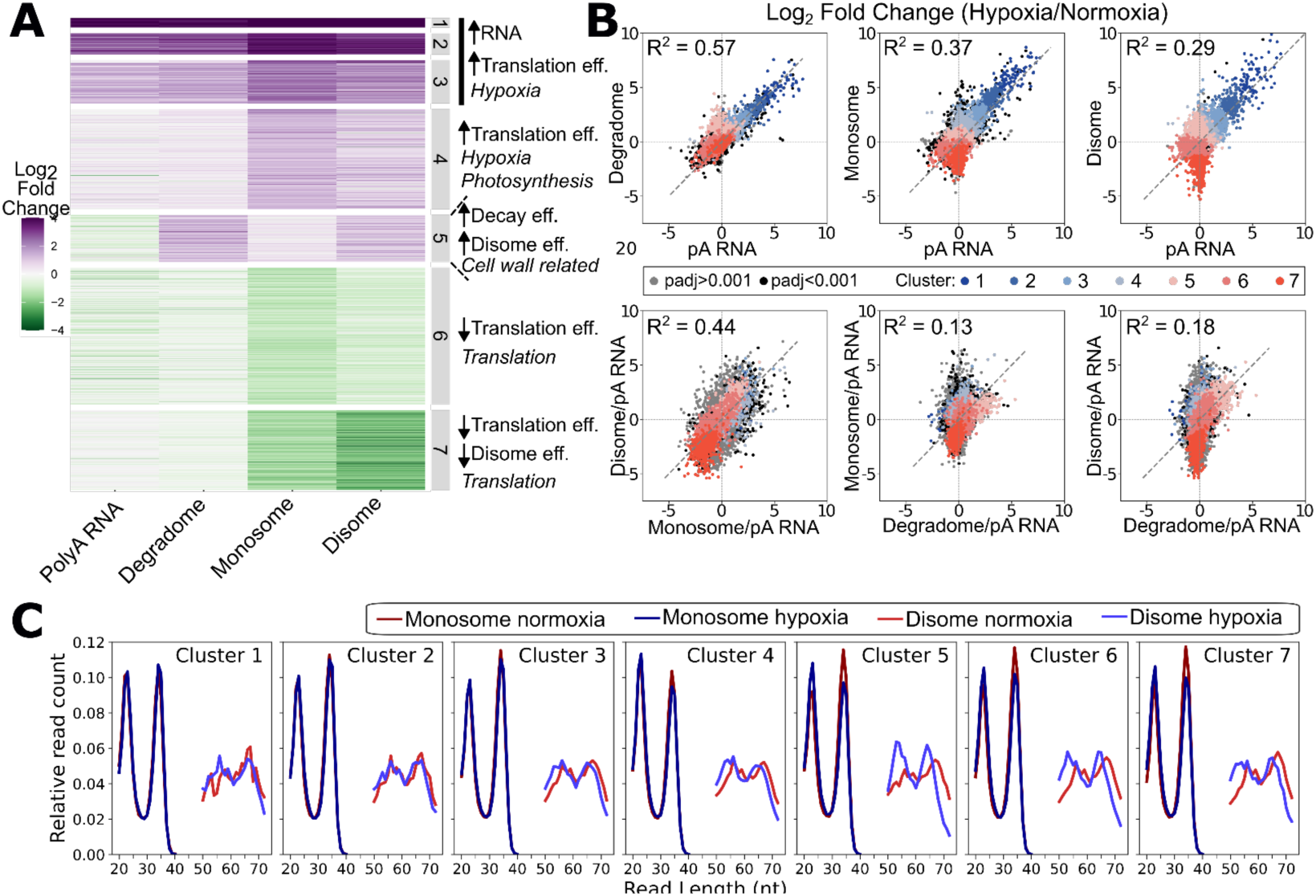
Translation coupled mRNA decay occurs under hypoxia for stress-induced mRNA and some transcripts associated with photosynthesis and growth. (***A***) Heatmap and clustering of similarly regulated genes under hypoxia stress. Genes with sufficient read counts in each data type and a significant difference between hypoxia and normoxia (|log_2_ FC| > 1; padj. < 0.01) in any of the four RNA data types are presented (3996 genes). Type of gene regulation and selected GO enrichment descriptors are indicated next to the clusters. GO enrichment results in ***SI Appendix,* Fig. S10** and **Dataset S2B**. (***B***) Correlation plots of Log_2_ FC (Hypoxia/Normoxia) of different data types (top row) or the Log_2_ FC of the calculated “efficiencies” (normalized against RNA abundance). (bottom row). Monosome/polyA (pA) RNA (MoEff) is commonly referred to as Translational Efficiency (TE) (57, 58). (***C***) Read length distribution of reads mapped to the transcripts in the individual clusters reveals that tightly stacked disomes coincide with increased co-translational mRNA decay in cluster 5.

Cluster 5 mRNAs stand out for a decrease in abundance that coincides with elevated disomes and 5’P reads. This indicates a conditional increase in decay coupled with disome formation. This cluster is enriched in genes related to the bioenergetically demanding processes of cell wall, lignin, and suberin biosynthesis (**Fig. S10**; **SI Dataset 2*B***). The high 5’P reads of cluster 5 mRNA contrasts with cluster 6 and 7 (**Fig. 3*A***; ***SI Appendix,* Fig. S11*A-B***). These show an insignificant change in mRNA and degradome reads, accompanied by similar or slightly greater reduction in monosome and disomes, indicating an overall reduction in translation.

Cluster 6 and 7 transcripts encode many proteins involved in photosynthesis and ribosome biogenesis. The regulation of ribosomal protein (RP) mRNAs seen here recapitulates prior observations in multiple species (45, 47, 56). The limited 5’ to 3’ decay of RP mRNA under hypoxia corresponds to their disengagement from polysomes, accumulation in UBP1 condensates (46), and nuclear retention (47). RP mRNAs re-engage with polysomes within minutes of reaeration, indicating these transcripts are safeguarded during the stress, by contrast to cluster 5 mRNAs.

Overall, total polyA mRNA levels correlate moderately well with degradome, monosome, and disome data, with R² values of 0.57, 0.37, and 0.29, respectively (**Fig. 3*B***). To discriminate transcriptional regulation from post-transcriptional regulation, read counts for each dataset type were normalized against the polyA RNA abundance for the same gene. For monosome-seq, this normalization is typically called translation efficiency (TE) (57, 58), but for clarity we refer to it as monosome efficiency (MoEff). Similarly, we define disome efficiency (DiEff) and decay efficiency (DegEff) for disome-seq and degradome-seq, respectively. MoEff and DiEff correlate reasonably well (R² = 0.44) (**Fig. 3*B***). This is unsurprising, since disome formation can be explained by two scenarios, (1) a stalled leading ribosome results in collision by a trailing ribosome, or (2) high rates of initiation of an mRNA cause ribosome crowding, elevating the chance of disome formation. The second scenario can explain the general correlation between MoEff to DiEff with the exception of some groups of genes as exemplified by cluster 5.

We find MoEff only weakly correlates with DegEff (R² = 0.13), whereas DiEff shows a slightly better correlation (R² = 0.18). Only a subset of DegEff data correlates with DiEff, largely attributable to transcripts in Cluster 5 (**Fig 3*B***). For transcripts of this cluster, the transcript levels decline upon hypoxia, concomitant with strong increases in degradome and disome reads.

In animals and fungi, ribosome stalling and the formation of collided disomes can induce NGD, which triggers turnover of transcripts through exogenous and endogenous cleavage events within the mRNA tunnel of the ribosome (18). Although NGD is described in the context of viral defense and a few other cases in plants (59, 60), ribosome stalling as a trigger is not well established. Cluster 5 transcripts show an increase in tightly stacked disomes under hypoxia, indicating that more tightly stacked or collided disomes promote co-translational decay (**Fig 3*C***). The increase in ribosome collisions on cluster 5 mRNAs could activate NGD, coupled with RaQC clearing of the nascent protein and recycling of ribosome subunits (20). Notably, mRNAs with a higher number of Asp codons do not exhibit a significant increase in disome reads under hypoxia despite the clear stalling of a subset (***SI Appendix,* Fig. S11*C-D***). A possible explanation is that localized increases in disome formation at a few Asp codons within a transcript may be obscured by the background level of disome reads.

### Conservation of ribosome stalling

Next, we sought to determine if peptidyl transferase center conformation (vacant versus occupied A-site), slow decoding of specific codons (start, stop, and amino acid), as well as uORF translation are conserved between Arabidopsis and a monocot model. To accomplish this, we profiled mono- and disome footprints of the shoot crown region harvested from V3-stage B73 maize. We found consistency in the ribosome footprint sizes corresponding to the two monosome and four disome types identified in Arabidopsis (**Fig. 4*A,B*; *SI Appendix,* Fig. S12**). The 3’ ends of footprints of vacant and occupied A-site ribosomes were 1 and 4 nts shorter, respectively, in maize than Arabidopsis. This could reflect differences in the efficiencies of RNase I cleavage or subtle structural distinctions between ribosomes of the two species.

**Fig. 4.**
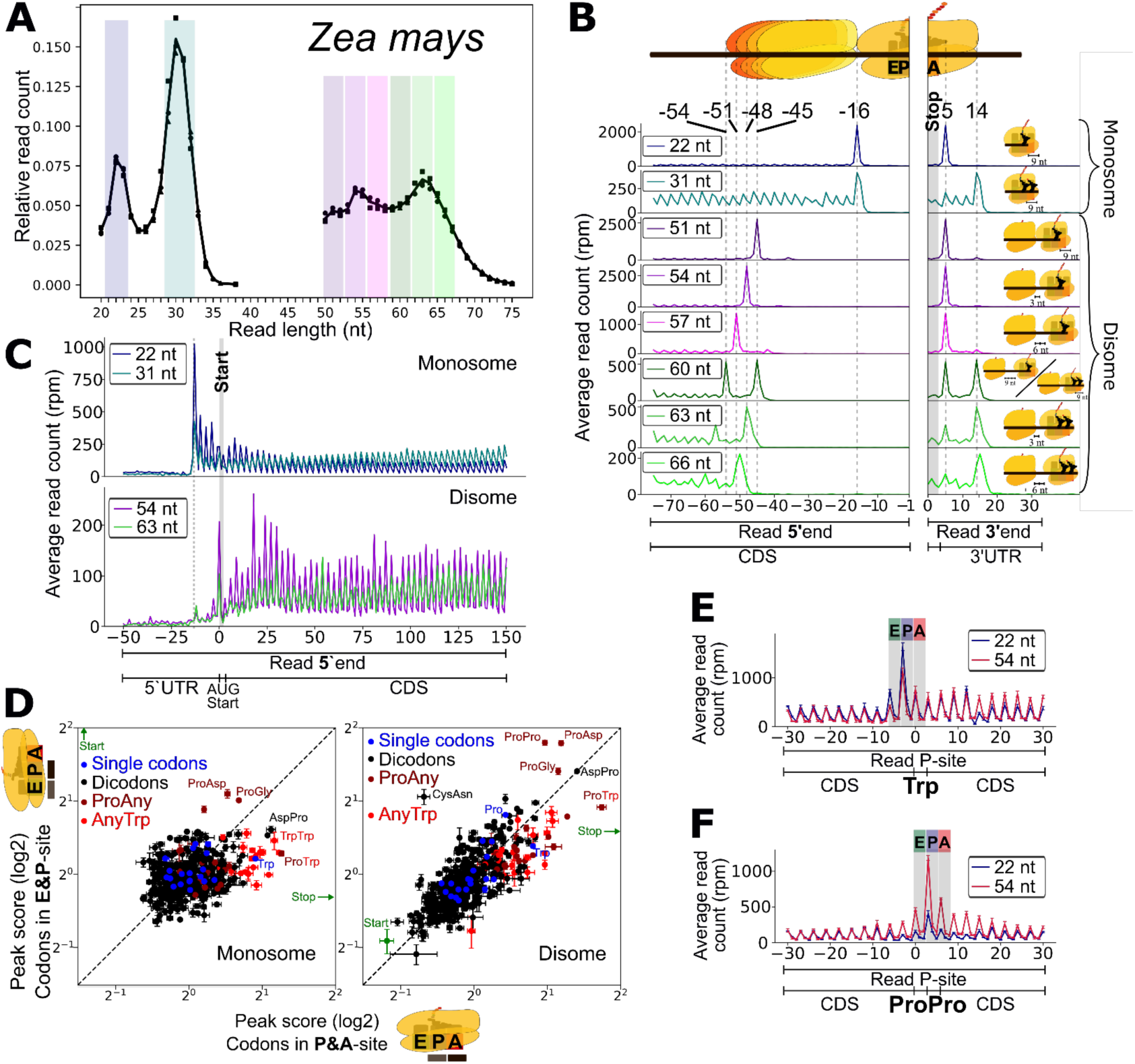
Disome pausing is prevalent at stop, di-Proline and Tryptophan codons in the shoot crown region of maize seedlings. (***A***) Read length distribution of mono- and disome-seq reads of maize shoot tissue. Lines represent averages; squares, dots, and triangles represent individual replicates. Colored vertical background bars indicate read lengths for the different ribosomal conformations per Fig. 1. (***B***) Average RPM of footprint 5’ or 3’-ends around the stop codon. (***C***) Average RPM of footprint 5’-end around the start codon and into the CDS. (***D***) Log_2_ peak score profiles of single amino acid (*y*-axis: P-site; *x*-axis: A-site) and dipeptide codons (*y*-axis: E&P site, *x*-axis: P&A site) for monosome and disome data. Start and stop codon peak scores outside the plotted range are indicated with arrows (input data: **Dataset S3**). (***E, F***) Average predicted P-site RPM distribution around single Trp (***E***) or ProPro (***F***) codons (***D, E, F***). Data and error bars in panels C-E represent standard deviation from 3 biological replicates. (***B,E,F***) *x*-axis units are distance from indicated codon in nt.

The maize monosome footprints peak at 22/23 and 30/31 nt, whereas disome footprints peak at 54 and 63 nt, representing 1-codon spaced ribosomes. The reduced footprint size of occupied A-site ribosomes led to a mixed representation of 60-nt disome footprints. These consist of both 3-nt spaced disomes with vacant A-sites and tightly stacked disomes with occupied A-sites, as indicated by the double peak at the stop codon (**Fig. 4*B*; *SI Appendix,* Fig. S12**). Even though both peaks are of similar height near the stop codon, we deduce that most 60-nt footprints represent tightly stacked A-site occupied ribosomes, since empty A-site ribosomes are overrepresented at the stop codon based on the monosome footprint data.

Similar to the translational ramp in Arabidopsis, maize monosomes transition from the empty A-site (22/23 nt) to occupied A-site form within the first 75 nt of the CDS seedlings (**Fig. 4*C***). Also consistent are the high peak scores observed for the initiation and termination codons when located in the P-site and A-site, respectively. Among codons, the Trp (UGG) exhibited the most prominent peak scores when occupying the A-site of the ribosome, suggesting its rate-limiting decoding in seedling tissue (**Fig. 4*D,E***). Pausing at Trp codons was not notable in Arabidopsis. Three di-codon motifs have high disome peak scores suggestive of stalling at di-Pro (E&P-site), Pro-Asp (E&P-site), and Pro-Trp (P&A-site) (**Fig. 4*D,F***).

We conclude that maize shares many mono and disome patterns with Arabidopsis, with subtle distinctions that could reflect tissue type and developmental stage or species-specific differences including codon usage or translational apparatus.

### Metabolite-induced stalling and CPuORFs

Next, we used our data to evaluate ribosome stalling events previously associated with metabolites and specific sequences within uORF or main (m)ORF coding regions. One such stall site called MTO1 is located within the mORF of *CYSTATHIONINE GAMMA-SYNTHASE 1* (*CGS1*) (61, 62). The 11 amino acid MTO region induces stalling under elevated concentrations of S-Adenosyl-L-methionine. We find MTO region stalling in our monosome and disome data that is coincident with a prominent 5’P peak 30 nt upstream of the stall site (***SI Appendix,* Fig. S7*F***). This validates MTO stalling and demonstrates it promotes mRNA cleavage just 5’ of the stalled ribosome.

Ribosome stalling on uORFs typically limits translation of the downstream protein-coding mORF. CPuORFs, identified in the 5’UTR of well over 100 Arabidopsis mRNAs, include some shown to cause ribosome stalling based on metabolite abundance (63). For many CPuORFs, the polypeptide sequence is conserved up to the stop codon. For some of these, ribosome stalling occurs near the stop codon but before polypeptide release (12, 50, 64–66). One hypothesis is that the stop codon may facilitate metabolite-mediated stalling due to the relatively slow kinetics of translation termination (67). To resolve uORF stalling, we surveyed CPuORFs undergoing translation in the Arabidopsis and maize datasets. In both species, monosome and disome footprints correspond to a paused leading ribosome with the termination codon positioned in the A-site of the ribosome on many CPuORFs (**Fig. 5*A*,*B***). Many termination stall sites are coincident with 5’P cleavage. These data confirm the hypothesis that unoccupied A-site ribosomes awaiting eRF1 at the the stop codon of CPuORFs is often coupled with co-translational decay.

**Fig. 5:**
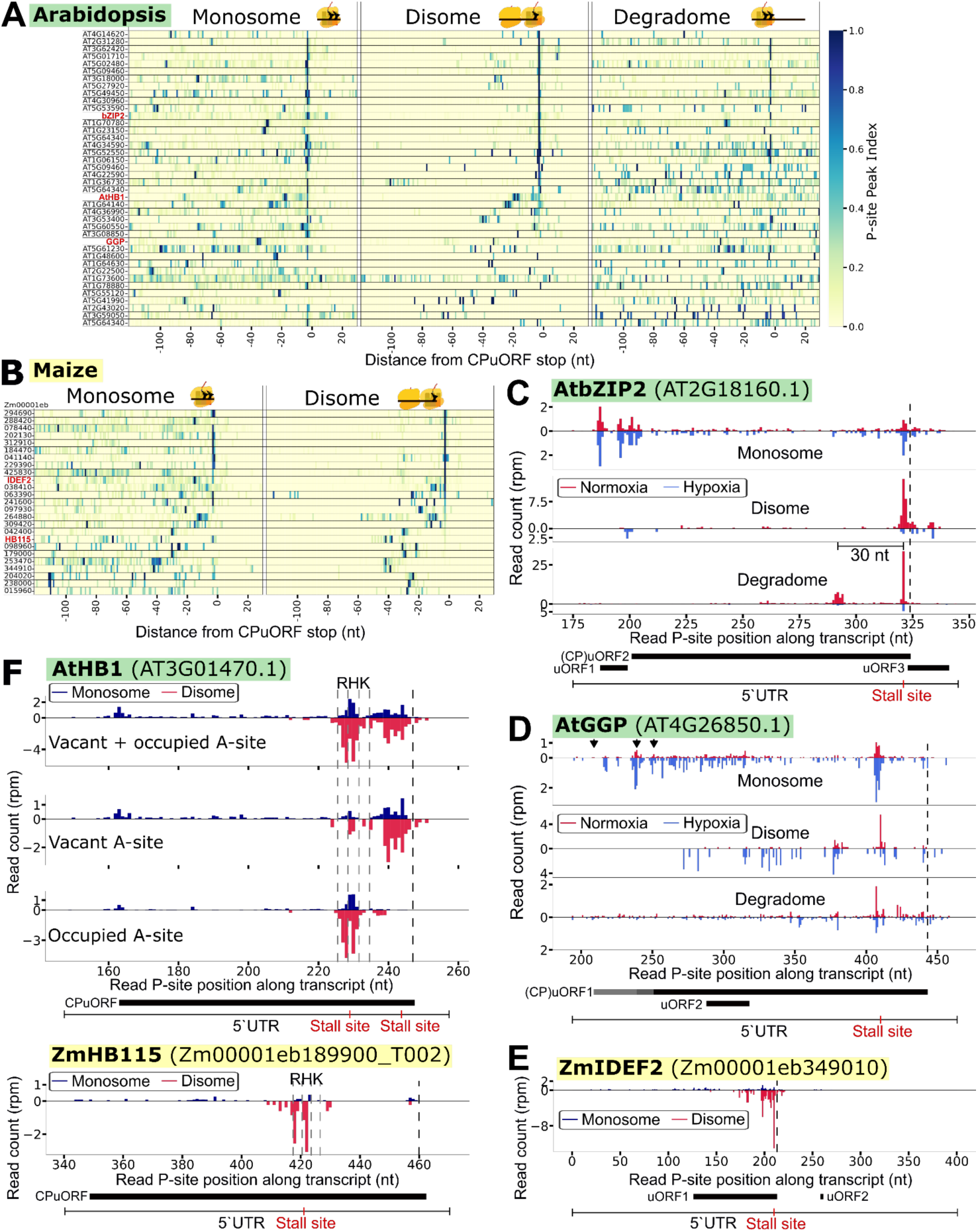
Mono- and disomes accumulate at the termination site of many CPuORFs represented in our Arabidopsis and maize datasets. (***A***) Arabidopsis (n=40) peak index of monosome, disome and degradome predicted P-site positions relative to CPuORF stop codon. *BASIC REGION/LEUCINE ZIPPER BOX2* (*bZIP2*) (AT2G18160) transcription factor, *ARABIDOPSIS THALIANA HOMEOBOX 1* (*AtHB1*, AT3G01470) and *GDP L-GALACTOSE PHOSPHORYLASE* (*GGP*/*VTC2,* AT4G26850) gene identifiers are highlighted in red. Genes shown had ≥ 5 disome reads within the CPuORF in all three biological replicates for normoxic seedlings. Zero (0) is the first nucleotide of the stop codon. (***B***) Peak index data for maize CPuORFs (n=20). *IRON-DEFICIENCY-RESPONSIVE ELEMENT BINDING FACTOR 2* (*ZmIDEF2,* Zm00001eb349010) and *HOMEOBOX DOMAIN 115* (*HB115, Zm00001eb189900_T002*) shown in red. (***C-F***) Average read count (RPM) at predicted P-site positions around *AtbZIP2* (***C***), *GGP* (***D***), *ZmIDEF2* (***E***), and *AtHB1* CPuORFs. For (***C***), (***D***) and (***E***) and the top panel of (***F***), all footprint sizes are used; in the middle panel of (***F***) only footprint sizes representing (leading) ribosomes with vacant A-sites were used; and for the bottom panel only footprint sizes representing (leading) ribosomes with occupied A-sites. (***D***) The *AtGGP* CPuORF does not contain an AUG start codon; potential non-AUG start codons are indicated by the downward black arrows.

Sucrose induces ribosome stalling at the stop codon of the CPuORFs of S1-group bZIP transcription factors (10), (11). Our Arabidopsis growth media included sucrose and ribosome stalling was evident at the characteristic location of S1-bZIP family members under normoxia (**Fig. 5*A,C***). Notably, disome accumulation and coincident 5’P peaks at these sites is reduced by hypoxia. The data identify other CPuORFs that induce ribosome stalling upstream of the stop codon, such as showrn for ascorbate-responsive CPuORFs of *GDP-L-GALACTOSE PHOSPHORYLASE* (*GGP*) (68) (**Fig. 5*A,D***). Recently, a translationally repressive and conserved uORF1 specific to monocots was identified in the *IRON-DEFICIENCY-RESPONSIVE ELEMENT BINDING FACTOR 2* (*OsIDEF2*) mRNA of rice (69), corresponding to *ZmIDEF2* of maize. Based on our data, monosomes occupy the length of uORF1 whereas disomes accumulate only 5’ of the stop codon (**Fig. 5*B, E***).

The codon-level examination of mono and disomes also provides insights for known but uncharacterized CPuORFs. For example, there is both an elongation and stop codon stall on the CPuORF of *HOMEOBOX 1* (*AtHB1*) (**Fig. 5*F***). This elongation arrest is characterized by mono- and disome footprint peaks of occupied A-site ribosomes six codons upstream of the stop codon with Arginine (Arg/R), histidine (His/H) and Lysine (Lys/K) codons in the E, P and A-site, respectively. This stalling is coincident with 5’P peaks (**Fig. 5*F***). In this unusual case, unoccupied A-site ribosomes peak at the stop codon without a 5’P peak. This pattern suggests that the ribosome peak at the stop codon is due to slow kinetics of translation termination or CPuORF-specific mechanisms. The latter interpretation is supported by the observation that the *AtHB1* CPuORF stop codon position is conserved in dicots, but not in monocots (70, 71). This raises the possibility of two distinct stall sites on this CPuORF: one broadly conserved, and a stop-codon-specific site present in dicots. Our maize data allow us to investigate these hypotheses. Stalling at the ArgHisLys (RHK) site of the CPuORF of *ZmHB115*, the *AtHB1* ortholog, indicates a conserved stalling mechanism at this site (**Fig. 5*F***). However, stalling was absent at the non-conserved stop codon position. Dicots may maintain two conserved stall sites, one at the conserved ArgHisLys site and another at the stop codon, whereas monocots could employ only the ArgHisLys stall. Our findings resolve the dual function of ribosome stalling on CPuORFs. These can both promote co-translational decay and limit mORF translation. The monitoring of ribosome conformation and proximity illustrates deep conservation and lineage specific innovations in CPuORF stalling signatures of plants.

## Conclusions

By integrating single codon resolution monosome-, disome-, and degradome-seq datasets, we resolve the position of ribosomes on mRNAs at high resolution, identifying monosomes with vacant or occupied A-sites and disomes that are tightly packed or spaced up to two codons. The data uncover multiple causes and characteristics of positional and conditional ribosome stalling that impact protein production and mRNA turnover (***SI Appendix,* Fig. S13*A, B***). We find ribosome stalling events associated with decoding of specific mono and di-codons can be deeply conserved, such as di-Pro pauses on *POLYUBIQUITIN* mRNAs associated with production of ubiquitin monomers or sites of disome pausing associated with a 5’P terminus (***SI Appendix,* Fig. S13*C***). The latter could occur when a trailing ribosome collides with a stalled ribosome, which is known to activate NGD as a surveillance mechanism in eukaryotes (3). Our discovery that collisions at Asp codons correlate with a reduction in cellular Asp content under hypoxia demonstrates conditional stalling and mRNA abundance control may be regulated by dynamics in amino acid pools (9).

This study provides new insight into the reorganization of the translational landscape by brief hypoxia (***SI Appendix,* Fig. S13*B***). First, we validate the high monosome translational efficiency of HRG mRNAs and the dissociation of RP mRNAs from translating ribosomes. Our results indicate the hypoxia-induced sequestration of RP mRNAs is uncoupled from 5’ to 3’ mRNA decay. Second, we find that conditionally collided disomes track high 5’P cleavage and reduced abundance of mRNAs required for energy demanding processes. Third, the data uncover the coupling of high translational efficiency (MoEff and DiEff) with high co-translational decay of HRG mRNAs. The high-flux of these coordinately-regulated mRNAs (47, 72), could facilitate their rapid clearing upon reoxygenation. Finally, the conditional pausing at Asp codons suggests amino acid levels can impact mRNA abundance and protein production.

Our comparative analysis of uORF translation in Arabidopsis and maize demonstrates that codon- and motif-specific stalls, and conditional uORF-mediated regulation can be broadly conserved. It is not yet clear whether the varied types of ribosome collisions observed (i.e., associated with unoccupied A sites including Asp codons under hypoxia and ribosomes just upstream of stall sites on CPuORFs) activate NGD. The coupling of Asp stalls with mRNA decay contrasts with di-Pro stalls that appear not to trigger decay of the proline-rich cell wall protein mRNAs, even under hypoxia. This difference may reflect the structural roadblock within the peptidyl transferase center during the formation of di-Pro peptide linkages (73). Hypusinated elongation factor 5A (eIF5A) specifically interacts with the P-site tRNA at di-Pro stalls to facilitate the necessary contortions. Neither the PPDQQ stalling motif on *POLYUBIQUITINs* nor other di-Pro codons in general are associated with 5’P monophosphate cleavage sites.

We hypothesize that ribosome collisions at single and combinations of codons activate a NGD pathway in plants. Plants encode orthologs of proteins necessary for NGD in mammals and yeast, with the notable exception of the Cue2 endonuclease that cleaves within the A-site of the collided ribosome to initiate the pathway (16, 21, 74). The plant process likely includes PELOTA/HBS1-mediated rescue of stalled ribosomes, coupled with 5’ decay via XRN4 and 3’ to 5’ decay involving the SUPERKILLER and exosome complexes. Plants also possess proteins of the RaQC pathway that pry apart the stalled ribosome, mark the nascent polypeptide for turnover, and recycle the stalled ribosome. In rice, RaQC proteins are crucial for pollen development and male fertility (75, 76). Based on this study, co-translational mRNA decay is triggered by closely spaced or collided ribosomes, but not in every circumstance. It remains to be determined if an endonuclease that cleaves between stalled ribosomes activates 5’ to 3’ decay or if decapping and XRN4 tethered to ribosomes are sufficient. We predict that protein synthesis and mRNA decay modulated by ribosome stalling, spacing, and A-site occupancy contribute to crucial changes in cell identity and state during the plant life cycle.

## MATERIALS AND METHODS

### Methods summary

*Arabidopsis thaliana* Col-0 seedlings were grown vertically on 0.5x MS medium under long-day conditions for 7 d. Hypoxia treatments were initiated at Zeitgerber time 16 by transferring plates to sealed chambers subjected to argon in darkness, and tissue was harvested after 2 h. Normoxic controls were maintained in matched chambers exposed to ambient air. *Zea mays* B73 plants were grown in soil under greenhouse conditions, and crown-region shoot base tissue was collected at the V3 stage.

To capture different ribosome states, frozen tissue was lysed in the presence of cycloheximide, digested with RNase I, and separated on sucrose gradients. Monosome and disome fractions were separated by sucrose gradient profiling, and ribosome-protected fragments of approximately 20–37 nt and 50–75 nt, respectively, were size-selected, and converted into indexed, UMI-containing libraries with Cas9-based depletion of abundant rRNA-derived products. In parallel, Arabidopsis poly(A) RNA-seq and 5′P/degradome-seq libraries were generated from the same samples to quantify transcript abundance and co-translational mRNA decay intermediates.

Reads were demultiplexed, adapter-trimmed, filtered against noncoding RNAs, deduplicated using UMIs, and aligned to the Arabidopsis TAIR10/Araport11 or maize B73v5 annotations. Aligned reads were used for downstream analyses.

Full growth conditions, library construction protocols, alignment parameters, filtering criteria, and custom analysis procedures are provided in the **SI Materials and Methods**.

## Data Availability

Raw sequencing data files and processed featurecount file, and read list (RiboWaltz output) files with predicted P-site positions per footprint are deposited at NCBI GEO with accession number GSE327520. Supplementary datasets provide processed data and comparative analyses.

## Author contributions

S.v.d.H., J.B.-S., and J.L.G designed research; S.v.d.H. performed research; S.v.d.H., J.B.-S., and J.L.G analyzed data; and S.v.d.H., J.B.-S., and J.L.G wrote the paper.

## Acknowledgements

This research was supported by an EMBO long-term postdoctoral fellowship grant (ALTF 264-2021) to S.v.d.H., the US National Science Foundation (MCB-1716913), MacArthur Foundation and USDA NIFA Hatch funding to J.B.-S, and the National Institute of General Medical Sciences of the National Institutes of Health (R35GM151048) and USDA NIFA Hatch 7002327 to J.L.G. We thank Mauricio Reynoso and Jeoffrey George for thoughtful discussions.

