## Supporting Information for "Ribosome stalling position, spacing, and A-site occupancy impact translation and co-translational mRNA decay in plants"

**This PDF file includes:**

Supporting
Supplemental Results
Figures S1 to S14
SI References

**Other supporting materials for this manuscript include the following:**

Datasets S1 to S5
Files S1 to S3

### MATERIALS AND METHODS

**Plant growth and treatment.** Arabidopsis (*Arabidopsis thaliana*, Col-0) seeds were sterilized by incubation in 70% (v/v) ethanol for 5 min, followed by incubation in 20% (v/v) bleach and 0.01% (v/v) Tween-20, and washed five times in double distilled water followed by dissolving the sterilized seeds in 0.01% (v/v) agar. Sterilized seeds were placed onto 0.5X Murashige & Skoog medium (0.5X MS salts, 0.4% [w/v] Phytoagar (PlantMedia, Dublin, OH), and 1% [w/v] Suc, pH 5.7-5.8) in 12x12 cm Petri dishes and stratified by incubation at 4°C for 3 d in complete darkness. After stratification, the plates were placed vertically into a growth chamber (Percival) with a 16-h light/8-h dark cycle at ~120 µmol photons$\cdot s^{-1} \cdot m^{-2}$ (cat. no. 21770; Sylvania), at 22°C for 7 d. For hypoxia stress, plates with seedlings were removed from the growth chamber at Zeitgeber time (ZT) 16 and subjected to HS by bubbling argon gas into sealed chambers in complete darkness for 2 h at room temperature. The chamber setup was as detailed previously (45). Oxygen partial pressure in the chamber was measured with a NeoFox Sport $O_2$ sensor and probe (Ocean Optics). Hypoxia ($O_2$ < 1.5%) was attained within 45 min of initiation of the stress. Control (normoxia) samples were placed in an identical chamber that was open to ambient air under the same light and temperature conditions. Tissue was rapidly harvested into liquid $N_2$, directly ground using a pestle and mortar and stored at -80°C. Three biological replicate experiments were performed on separate days.

Corn (*Zea mays*, inbred line B73) seeds were rinsed with cold tap water and imbibed in 40 mL water overnight. Surface sterilization was performed by immersing seeds in 40 mL of 5% [w/v] bleach for 15 min with occasional mixing. Seeds were then incubated on two layers of filter paper in darkness at 28°C for 2 d. Germinated seeds were transferred to soil (Sungro Sunshine Mix #5 supplemented with Osmocote and Marathon) and grown in a greenhouse. Plants were watered three times per week, with one watering supplemented with chelated DTPA iron (330 Fe). At the V3 stage (~14 d post-transfer to soil), the base of the stem (crown region) was harvested at solar noon. This region includes the shoot apex, leaf primordia, leaf base and stem tissue. To do so, roots were removed, followed by excision of a 1 cm section from the basal stem. Shoot base sections from four plants were pooled per biological replicate.

**RNase I digestion test and diagnostic polysome profiles.** Frozen pulverized tissue was lysed and RNAse I treated as described previously (77), with minor modifications. In brief, ~200 µl pulverized Arabidopsis or corn tissue was mixed with 600 µl ice cold extraction buffer [100 mM Tris-HCl (pH 8.0), 40 mM KCl, 20 mM $MgCl_2$, 2% (v/v) polyoxyethylene (10) tridecyl ether, 1% (w/v) deoxycholic acid, 1 mM DTT, 0.1 mg/mL cycloheximide, 10 U/ml DNase I], briefly vortexed and incubated on ice for 10 min, followed by centrifugation at 16,000 x *g* for 15 min at 4°C. The supernatant was transferred to a new tube and 200 µg (~200 µl) was incubated with different concentrations of RNase I (Ambion, AM2294) and incubated for 2 h at root temperature or 5 µl SuperaseIn RNase inhibitor (Ambion, AM2696) was added as control and the sample put on ice. Using our established method (78), lysates were loaded onto 4.5 mL precooled 15-50% (w/v]) sucrose gradients [40 mM Tris·HCl (pH 8.4), 20 mM KCl, 10 mM $MgCl_2$, and 5 µg/mL cycloheximide] and spun at 4ºC for 90 min at 50,000 rpm in a Beckman LM centrifuge using a SW55.1 rotor. Gradients were analyzed using a Brandel SYN-202 Density Gradient Fractionator and Detector system, connected to a chart recorder and fraction collector. Samples were placed immediately on ice.

**Monosome- and disome-seq libraries.** Ground Arabidopsis material was lysed, RNAse I treated, and sucrose gradient fractionated as described above with the exception that the RNase I treatment was performed with 0.5 U/µg RNA in 400 µl for 1 h at room temperature. An RNase I concentration of 0.5 U/µg RNA was used as concentrations above 1U/µg RNA compromised ribosome integrity (***SI Appendix,*** **Fig. S1**). Based on the UV profile, three to four fractions of ~100 µl each corresponding to monosomes and disomes were collected from the sucrose gradients. RNA was purified from these samples according to the TRIzol reagent manual. Briefly, 900 µl TRIzol (ThermoFisher Scientific) and 200 µl chloroform were added and a 30 min precipitation was following addition of 950 µl isopropanol, 35 µl NaOAc and 4 µl glycogen (5 mg/mL ThermoFisher) at -80°C, followed by an 80% (v/v) ethanol wash. The final pellet was dissolved in 10 µl 10 mM Tris-HCl (pH 8.0).

RNA footprint library preparation was performed as described (79), with some modifications. In brief, the RNA was run on a 15% (v/v) Urea polyacrylamide gel [1X TBE, 48% (w/v) Urea, 0.08% (w/v) ammonium persulfate, 0.2% (v/v) N,N,N',N'-Tetramethylethylenediamine], stained with SYBR Gold™ (Invitrogen) and bands between ~20-37 nt and ~50-75 nt were extracted for monosomes and disomes footprints, respectively. The gel slices were incubated in RNA gel extraction buffer [300 mM sodium acetate, 1 mM EDTA and 0.25% (w/v) SDS] overnight at room temperature with gentle agitation. RNA was extracted using isopropanol and glycogen, dephosphorylated using T4 Phosphonucleotide kinase and linker ligated (79) using linkers containing 5 nt unique molecular identifiers (UMI) and 5 nt indexes. RNA fractions corresponding to linker ligated ribosome footprints were cut from 10% (w/v) Urea-polyacrylimide gels and extracted overnight in the RNA gel extraction buffer following precipitation. Up to six samples with different linker indexes were pooled into 10 µl MiliQ $H_2O$ and used for cDNA synthesis using Protoscript II (New England Biolabs) using a RT primer that contains another 2 nt UMI (79), following hydrolysis of the RNA using high NaOH and cleanup with the oligo Clean & concentrator kit (Zymo research) eluting in 8 µl water. cDNA was run on 10% (w/v) Urea-polyacrylamide gel and appropriate cDNA bands were extracted from the gel and cDNA was extracted using DNA gel extraction buffer [300 mM NaCl, 10 mM Tris-HCl (pH 8.0), 1 mM EDTA] and precipitated; finally the cDNA was circularized using CircLigase II ssDNA ligase (BioResearch Technologies, CL9021K) (79). See **Dataset S4A** for all linker, UMI, indexes and barcode info.

Circularized cDNA was amplified using Phusion polymerase and appropriate barcoded primers (**Dataset S4A**) and treated with Cas9 using a variety of gRNAs to remove rRNA fragments. For Cas9 treatment; gRNA targets were identified using Ribocutter software (80) and synthesized using EnGen® sgRNA Synthesis Kit, *S. pyogenes* (NEB, E3322V) (**Dataset S4B**). Next, 1.5 µl of 20 µM SpCas9 (NEB, M0386T) was mixed with 3 µl 34.5 µM gRNA mix (total MW), 6.5 µl 10X Cas9 reaction buffer and 4 µl water and incubated for 10 minutes at 25°C. 50 µl of the Phusion PCR reaction mixture was then mixed with 15 µl of the Cas9-gRNA mix and incubated for 1 h at 37°C, followed by heat inactivation for 5 min at 65°C and removal of the gRNA with 1.5 µl 10 U/µl RNase I (Ambion) at 37°C for 10 min. The library was then purified using 1.5X Ampure XP beads according to manufacturer instructions, resuspended in 20 µl 10 mM Tris-HCl (pH 8.0). The cleaned library was reamplified by Phusion PCR using "EnrichS1" and "EnrichS2" primers (**Dataset S4A**), run on an 8% (w/v) TBE polyacrylamide gel and bands from the stated size range were excised from the gel and DNA was extracted in DNA gel extraction buffer and precipitation, and finally dissolved in 15 µl 10 mM Tris-HCl (pH 8.0) as the final library. All libraries were sequenced with 150 nt paired-end reads by Novogene, but only the forward reads (read 1) were used for analyses.

**PolyA RNA- and degradome-seq libraries.** *PolyA RNA isolation and sequencing.* Total RNA was extracted from approximately 100 µL pulverized Arabidopsis seedling tissue using TRIzol reagent. After phase separation with chloroform, 500 µL of the aqueous phase was mixed with 500 µL ethanol and RNA was purified using Zymo-Spin IICR columns (RNA Clean & Concentrator-25 kit, Zymo Research). On-column DNase I treatment was performed according to the manufacturer's instructions. Poly(A) RNA was isolated from 25 µg total RNA diluted in 200 µL binding buffer (100 mM Tris-HCl pH 8.0, 1 M LiCl, 10 mM EDTA pH 8.0, 1% (w/v) SDS, 5 mM DTT, 1.5% Antifoam A). One µL of 12.5 µM 20-nt biotinylated oligo(dT) was added and incubated for 10 min at room temperature. Poly(A) RNA was captured using 20 µL streptavidin-coated Dynabeads (M-280; Thermo Fisher Scientific). Beads were washed sequentially with buffer A (10 mM Tris-HCl pH 8.0, 150 mM LiCl, 1 mM EDTA, 0.1% SDS), buffer B (10 mM Tris-HCl pH 8.0, 150 mM LiCl, 1 mM EDTA), and low-salt buffer. Poly(A) RNA was eluted at 80°C for 2 min in 25 µL elution buffer (10 mM Tris-HCl pH 8.0, 1 mM EDTA). The polyA RNA was used for both polyA RNA-seq and degradome seq. Poly(A) RNA-seq libraries were prepared as described (81).

*Degradome-seq library.* Degradome-seq (5'P) libraries were prepared in a similar fashion as described (26, 48, 82). Briefly, 5 pmol single-stranded RNA linker (**Dataset S4A**) was ligated to 100–400 ng poly(A) RNA using T4 RNA ligase 1 (NEB). The excess linker was removed by an additional round of poly(A) RNA purification. RNA was purified using 1.8X RNAClean XP beads (Beckman Coulter) and eluted in 11 µL water. Reverse transcription was performed using 1.3 µL 20 µM 5P-seq-RT primer in a 20 µL SuperScript IV reaction (Thermo Fisher Scientific). RNA was hydrolyzed by incubation with 0.1 M NaOH at 65°C for 20 min and neutralized with 0.1 M Tris-HCl (pH 7.0), followed by cleanup using 1.8X AMPure XP beads. cDNA was amplified for six cycles, treated with Cas9 using three sgRNAs, and further amplified as described for monosome- and disome-seq libraries, using degradome-specific barcoded primers (**Dataset S4**). Final PCR products were purified using a two-step AMPure XP bead cleanup (0.5X followed by 0.4X), washed twice with ethanol, and eluted in 10 µL 10 mM Tris-HCl (pH 8.0).

**Computational processing and alignment of sequencing data.** *Mono- and disome-seq alignment (Arabidopsis thaliana).* Reads were split per sample based on the linker index and linker index and adapter trimmed using cutadapt (v4.1) using sequence 5'-iiiiiAGATCGGAAGAGCAC (i=linker index). Reads shorter than 10 nts after trimming were discarded. Trimmed reads were aligned to non-coding (nc)RNAs, consisting primarily of rRNA and tRNA (**File S2**), using bowtie2 with the following parameters: --local -L 20 --score-min G,9,8 trim5 2 trim3 5. Unmatched reads were used for further analysis. Duplicated reads were removed using the unique molecular identifier (UMI) made up by the 2 and 5 nt random sequences surrounding the footprint, with a custom Python script and then the UMI was removed. The remaining reads were aligned to the *Arabidopsis thaliana* TAIR10 reference genome and Araport11 annotation (Ensembl release 36) using STAR (v2.7.11a) with the following settings: --alignIntronMax 5000 --alignIntronMin 15 --outFilterMismatchNmax $mismatch --outFilterMultimapNmax 20 --outFilterType BySJout --alignSJoverhangMin 8 --alignSJDBoverhangMin 2 --outSAMtype BAM SortedByCoordinate --quantMode TranscriptomeSAM --outSAMmultNmax 1 --outMultimapperOrder Random, where $mismatch was set to 1 for monosome-seq and 2 for disome-seq datasets.

Maize (*Zea mays*) mono- and disome-seq data were processed analogously, using a maize-specific ncRNA reference (**File S3**) and the Zm-B73-REFERENCE-NAM-5.0 genome assembly with its corresponding annotation.

*Degradome-seq alignment.* Degradome-seq reads were adapter-trimmed using cutadapt (v4.1) using adapter sequence 5'-AGATCGGAAGAGCACACGTCTGAACTCCAGTC-3'. Reads shorter than 65 nts after trimming were discarded. Next, reads were aligned to ncRNA using bowtie2 with –trim 8 option, unaligned reads were retained. UMIs were extracted using umi-tools, and aligned to the Arabidopsis TAIR10 genome and Araport11 annotation file (Ensembl release 36) using STAR (v2.7.11a) with the following parameters: --outSAMtype BAM SortedByCoordinate --alignEndsType Extend5pOfRead1 --outFilterMatchNminOverLread 0.9 --outSAMmultNmax 1 --alignIntronMax 5000 --outReadsUnmapped Fastx --limitBAMsortRAM 8000000000 --quantMode TranscriptomeSAM --outSAMheaderHD @HD VN:1.4. Aligned reads were deduplicated using umi_tools.

*PolyA RNA-seq alignment.* PolyA RNA-seq reads were adapter trimmed using cutadapt (v4.1) using the same adapter sequence as for degradome-seq. The first 8 nts were removed and reads shorter than 65 nts were discarded. Next, reads were aligned to ncRNA using bowtie2, unaligned reads were used for further analysis. Alignment to the genome was performed the same as for degradome-seq except that --alignEndsType was set to Local.

**Computational analysis of aligned reads.** Trimmed read lengths and 5` and 3` end positions were extracted from transcriptome alignment files (bam) using riboWaltz (83), together with annotated start and stop codon positions. Downstream analyses were performed using only representative gene models as defined by Araport11 for *Arabidopsis* and the canonical gene models defined by Zm-B73-REFERENCE-NAM-5.0-Zm00001eb for maize. Mitochondrial and chloroplast transcripts were excluded from all analyses, unless mentioned otherwise.

Distributions of 5′ and 3′ read ends around annotated start and stop codons were generated using a custom Python script (**Fig. 1C,D**). For each read length, the offset used to infer the ribosomal P-site position was defined as the nucleotide position corresponding to the highest reads per million (RPM) value upstream of the start codon (monosome-seq) or stop codon (disome-seq). For degradome data, the predicted P-site is assigned 13 nts downstream of the read's 5' end. This position was assigned as the first nucleotide of the P-site codon. Predicted P-site positions were used for all downstream analyses, including read-length distributions across 5′ UTRs, coding sequences (CDS), and 3′ UTRs (**Fig. 1*B* and 5*A*; *SI Appendix,* Fig. S3**). AT2G01021.1 and AT2G20410.1 were excluded from the read length distribution analysis, as these caused non-biological biases at specific read lengths.

*Peak score calculation.* Peak scores for codons, dicodons, and tricodons were calculated by dividing the number of reads at the inferred A-, P-, or E-site by the average read density of the surrounding region. The surrounding region comprised 14 codons upstream and 14 codons downstream of the codon(s) of interest, excluding the two codons immediately adjacent to the codon of interest. Only the first nucleotide of each codon was used, corresponding to the peak of the 3-nt periodicity. Log2-transformed peak scores were used for visualization and downstream analyses (**Fig. 2*A* and 5*C*; *SI Appendix,* Fig. S5; Datasets S2 and S3**).

*Signal peptide.* Signal peptide and subcellular localization predictions generated independently by AtSubP (84) and TargetP (85) were downloaded from TAIR (March 9, 2024). For each prediction tool, transcripts were grouped according to the localization categories shown in the corresponding figure. RPM values at predicted P-site positions were normalized within each transcript by dividing the RPM at each position by the total P-site RPM for that transcript. Normalized profiles were then averaged across all transcripts within each localization group to generate the final positional distributions (***SI Appendix,* Fig. S2**).

*Probability plots.* Sequence features surrounding predicted ribosomal stall sites were analyzed using kpLogo (86). Only stall sites located within CDS regions were considered. For each site, a window spanning 30 codons upstream and 9 codons downstream of the P-site codon was extracted. Sites were classified as stall sites if they contained at least five ribosome footprint reads with predicted P-sites at that position and exhibited a ≥20-fold enrichment relative to the surrounding region, calculated as described for peak scores. These criteria had to be met in all three biological replicates. kpLogo was run with a Bonferroni-adjusted positional p-value cutoff of 0.01 (-pc 0.01). Input FASTA files are provided in **File S1**.

*Fast fourier transformation (FFT) signal calculation.* FFT signals were calculated to determine nt periodicity in degradome data, and was calculated using the fivepseq package (87).

*CPuORFs.* Stop codon positions for Arabidopsis CPuORFs used in **Fig. 5A** were extracted from van der Horst et al., 2019 (63). Maize CPuORFs were identified by first extracting all ORFs from B73v5 cDNA and translating them to protein sequences using the EMBOSS getorf package (https://www.bioinformatics.nl/cgi-bin/emboss/help/getorf) (88). ORFs with exact matches to Zea mays annotated proteins were removed, only leaving ORFs outside the annotated coding regions. Arabidopsis CPuORF sequences were extracted from https://theeukaryoticcpuorfdatabase.github.io/arabidopsis.html; (*Oryza sativa*) CPuORF sequences were extracted from Takahashi et al., 2020 (89). Both Arabidopsis and rice CPuORF sequences were aligned against the remaining maize ORF amino acid sequences using BLASTp. Next, non-homologous transcripts were removed using the ensembl-compara homology file (release 62) for *Zea mays*. Identified transcript with the orthologous CPuORFs and their stop codon positions used for **Fig. 5A,B** can be found in **Dataset S5**.

### Supplemental Results and Methods

#### Ribosome stalling and translational control observed on additional gene groups

*Ribosome stalling on ubiquitin-ribosomal protein fusions*

Two families of eukaryotic ribosomal protein have a N-terminal ubiquitin monomer (*RPL40/eL40; RPS27a/S31Y*). Our data provide evidence of ribosome stalling at the PPDQQ on both *eL40* and two of the three *S31Y* gene family member mRNAs, but within the ubiquitin monomer sequence (***SI Appendix,*** **Fig. S6*D***). In yeast, the processing of the N-terminal UBI fusion on eL40 and eS31 is required for ribosome function, and is suggested to occur very rapidly and likely co-translationally, as the intact precursor protein is not detected by standard methods (90). By contrast, in mammals, the removal of the N-terminal UBI on eS30 is carried out by a nucleolar deubiqutinase (DUBs) during ribosome biogenesis (91). Given that ribosome stalling on these RPs in Arabidopsis is within the ubiquitin monomer sequence rather than near the site where the DUB would clear, we propose the pausing is unrelated to co-translational cleavage.

#### Reduced MoEff and DiEff and increased DegEff on transposable element RNAs

Transposable element RNAs are reportedly less efficiently translated, enriched with disomes, and more prone to decay in rice callus and Arabidopsis *ddm1* (*DNA METHYLATION 1*) mutants (92). *ddm1* plants display elevated transposable element expression. To determine if similar patterns are observed in our data, 48 transposable element RNAs with sufficient expression were analyzed (see Methods below). We observed a slight reduction in MoEff and an increase in DegEff, similar toKim et al. (92) but contrary to their findings, DiEff was reduced (***SI Appendix,*** **Fig. S14**). This discrepancy may be attributed to differences in the experimental conditions or their use of a mutant with high transposable element expression.

### Methods

*Transposable elements methods.* Transposable elements were analyzed using the same alignment strategy described above, but with an annotation file that included transposable elements (Araport11_GTF_genes_transposons.Jan2024.gtf). Read counts were obtained using featureCounts using the following parameters "featureCounts -T 4 -t CDS,transposable_element_gene -g gene_id -O -s 1 -J -G". Transcripts per million (TPM) was calculated (((number of reads / transposable element length)/ total number of reads) x 1,000,000) for RNA-seq, monosome-seq and disome-seq. For degradome-seq data, Reads per million (RPM) ((number of reads / total number of reads) x 1,000,000) were used, as only one fragment per transcript can be detected.

*Differential gene expression.* For differential expression analyses, CDS-level read counts were extracted using featureCounts (Subread v2.0.6) with the parameters: -T 3 -t CDS -g gene_id -O -s 1 -J -R CORE -G. Count matrices were analyzed using deltaTE, which also calculated "efficiencies" (MoEff, DiEff, and DegEff) (58). Resulting log2 fold changes and adjusted p-values were used for **Fig. 3; *SI Appendix,* Fig. S10** and are provided in **Dataset S1A.** Genes with adjusted p-values < 0.01 and a log2 fold change below -1 or above 1 in at least 1 seq type (pA RNA, degradome, monosome or disome-seq) with sufficient sequencing depth across all datasets, were retained, resulting in 3995 transcripts. These transcripts were mapped in 7 clusters and for each cluster GO enrichments were calculated and visualized with clusterProfiler (93).

**Fig. S1.** Digestion of polysomes with high concentrations of RNAse I reduces 80S monosome stability.

**Fig. S2.** Degradome data show 3-nt periodicity, particularly in hypoxia treated samples.

**Fig. S3.** No indication of ribosome stalling near the start codon of mRNAs with signal peptides.

**Fig. S4.** Read length distribution

**Fig. S5**. P-site RPM distribution around individual Asp and Pro codons.

**Fig. S6**. $Log_2$ peak score profiles of dipeptide codons comparing normoxia and hypoxia.

**Fig. S7.** Average read counts (RPM) at predicted ribosomal P-site positions across different transcripts.

**Fig. S8.** Hypotheses for explaining different ribosome stalling events.

**Fig. S9.** Probability logos of amino acid and mRNA sequences at ribosomal pause sites identified in degradome, monosome, and disome datasets of Arabidopsis.

**Fig. S10.** Gene ontology analysis of different clusters in **Fig. 3**.

**Fig. S11.** Correlation plots of Log2 FC (Hypoxia/Normoxia) across different data types.

**Fig. S12.** Monosome and disome footprint distribution around the start and stop codon of shoots from vegetative stage V3 B73 maize dataset.

**Fig. S13.** Summary model for this study.

**Fig. S14.** Transposable element transcripts are less efficiently translated and more prone to degradation.

**Dataset S1.** Raw codon pause score of single amino acids, dipeptides and tripeptides for *Arabidopsis thaliana* data used for **Fig. 2*A*** and **S6**.

**Dataset S2.** deltaTE output, GO enrichment results and raw TPM or RPM data.

**Dataset S3.** Raw codon pause score of single amino acids and dipeptides for *Zea Mays* data used for **Fig. 4*D***.

**Dataset S4.** Lists of RNA and DNA oligos used in this study.

**Dataset S5.** Full transcript names and associates stop codon position used for **Fig. 5*A,B***.

**File S1.** Zip file containing fasta files used as input for creating the probability plots in **Fig. 2*D*** and ***SI Appendix,* Fig. S9**.

**File S2.** Fasta file containing sequence list used to remove rRNA and tRNA from *Arabidopsis thaliana* read files.

**File S3**. FASTA file containing sequence list used to remove rRNA and tRNA from *Zea mays* read files.

### Supplemental Figures

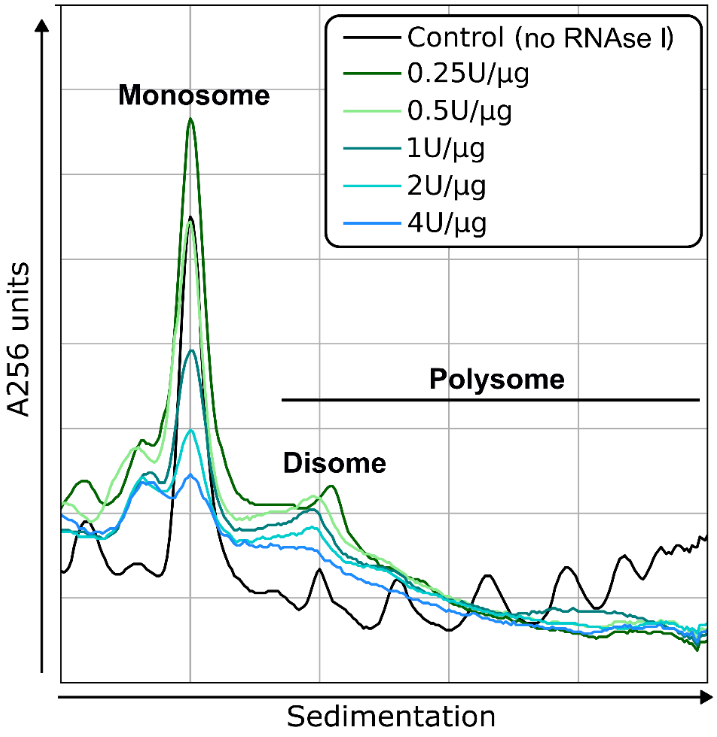


**Fig. S1.** Digestion of polysomes with high concentrations of RNAse I reduces 80S monosome stability. Representative UV absorbance (256 nm) profiles of polysomes separated by sedimentation through 15-55% (v/v) sucrose density gradients following digestion with increasing units of RNase I per microgram RNA. The largest peak corresponds to 80S monosomes.

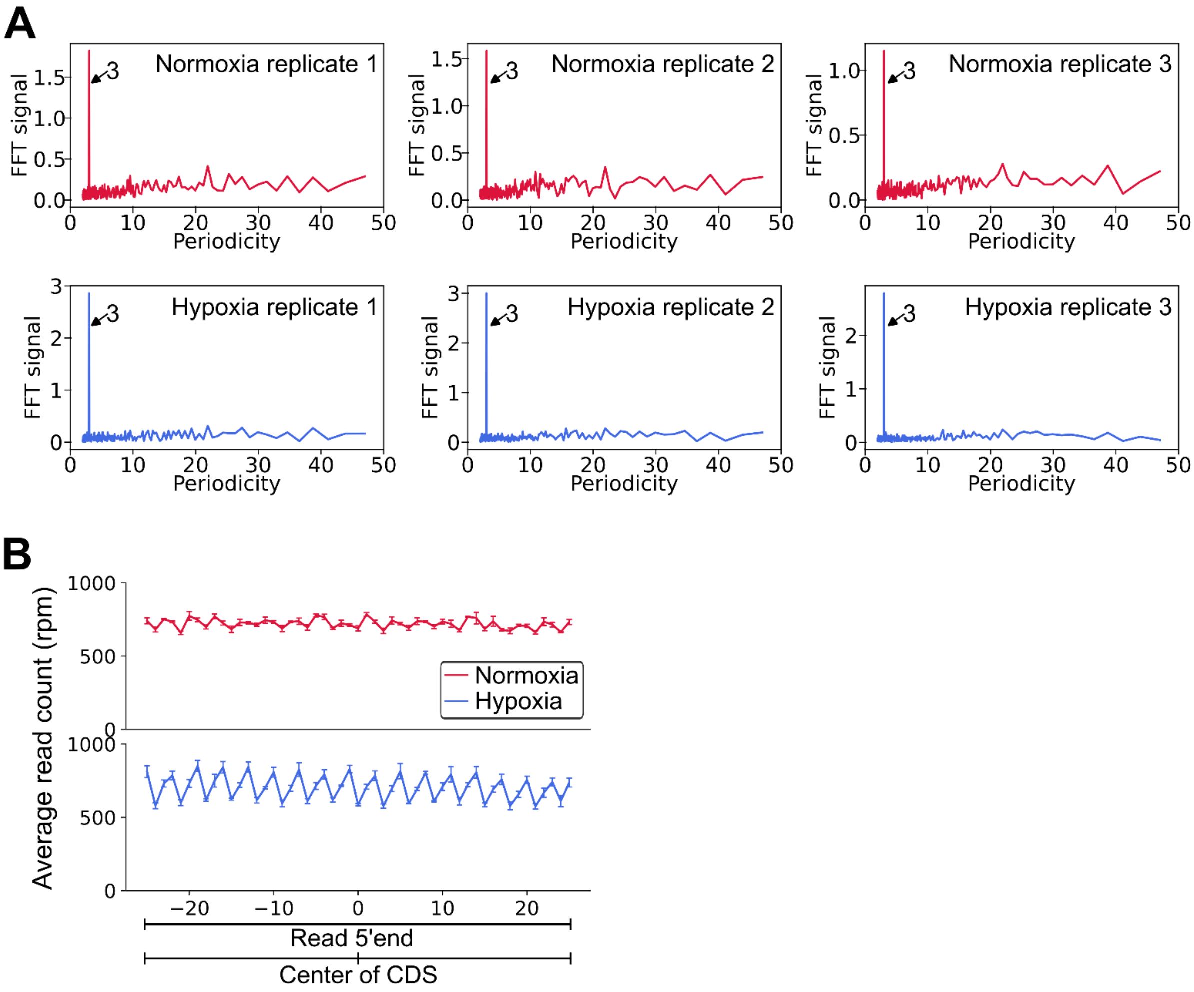


**Fig. S2.** Degradome data show 3-nt periodicity, particularly in hypoxia treated samples. (***A***) Fast Fourier Transformation (FFT) signals near the start codon, calculated using 5'P degradome-seq, show strong 3-nt periodicity. (***B***) Read 5' end distribution around the center of the coding sequence of all transcripts. Error bars represent standard deviation from three biological replicates.

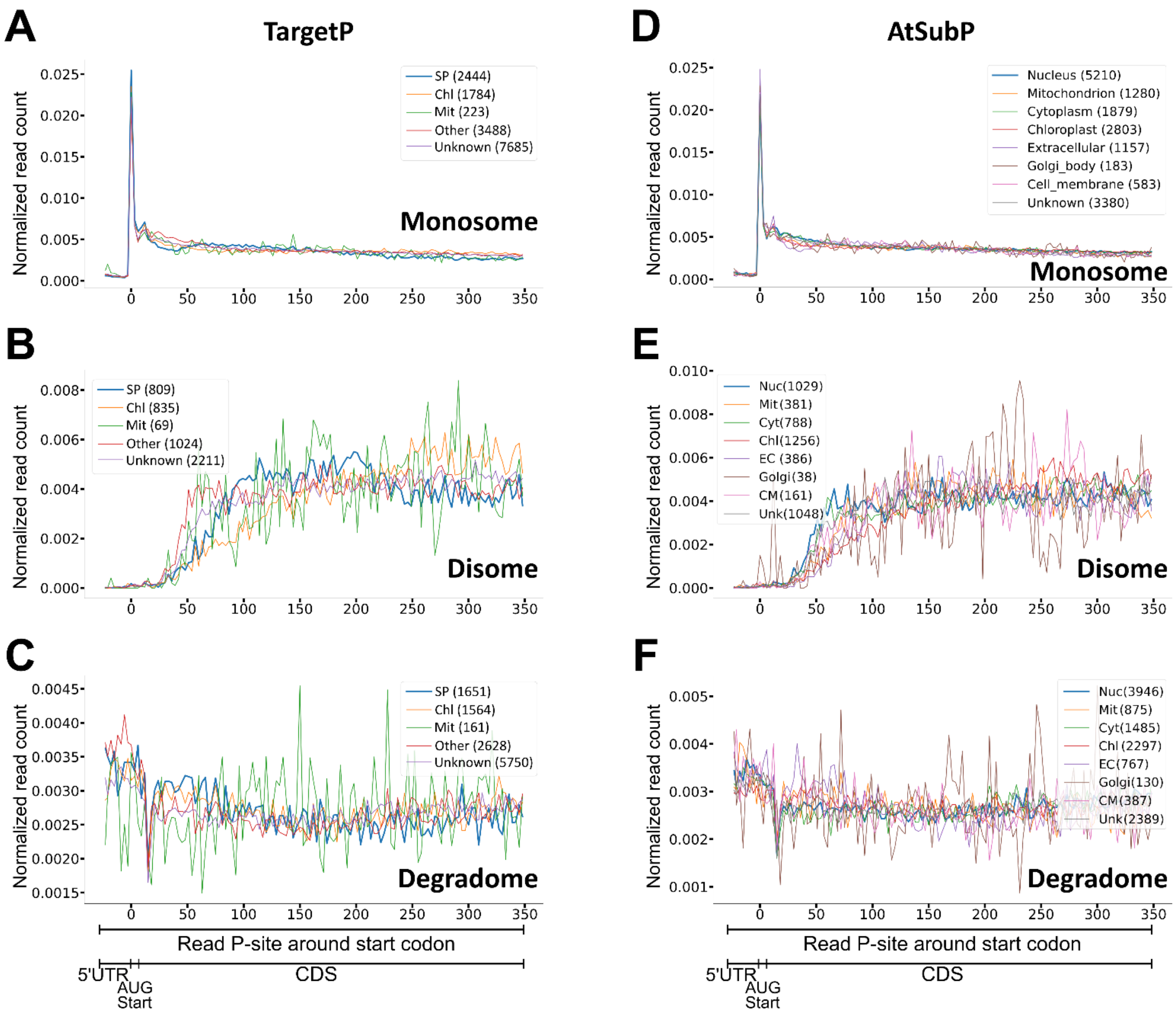


**Fig. S3**. No clear indication of ribosome stalling near the start codon of mRNAs with signal peptides. TargetP and AtSubP database predictions were used to map monosome, disome and degradome reads proximal to the start codon region (first 350 nucleotides of CDS) of candidate genes. The number of reads (monosome, disome, degradome) was counted for each predicted ribosome P-site position and then normalized within each transcript by dividing the reads at a given P-site by the total number of reads for that transcript. Next, transcripts were grouped based on their predicted localization, as predicted by TargetP (***A-C***) or AtSubP (***D-F***). The normalized read counts were averaged across transcript product localization groups. Only the first nt of each codon is shown for clarity and hence no codon periodicity is visible. TargetP abbreviations are: SP: secretory path, Chl: chloroplast, Mit: mitochondria.

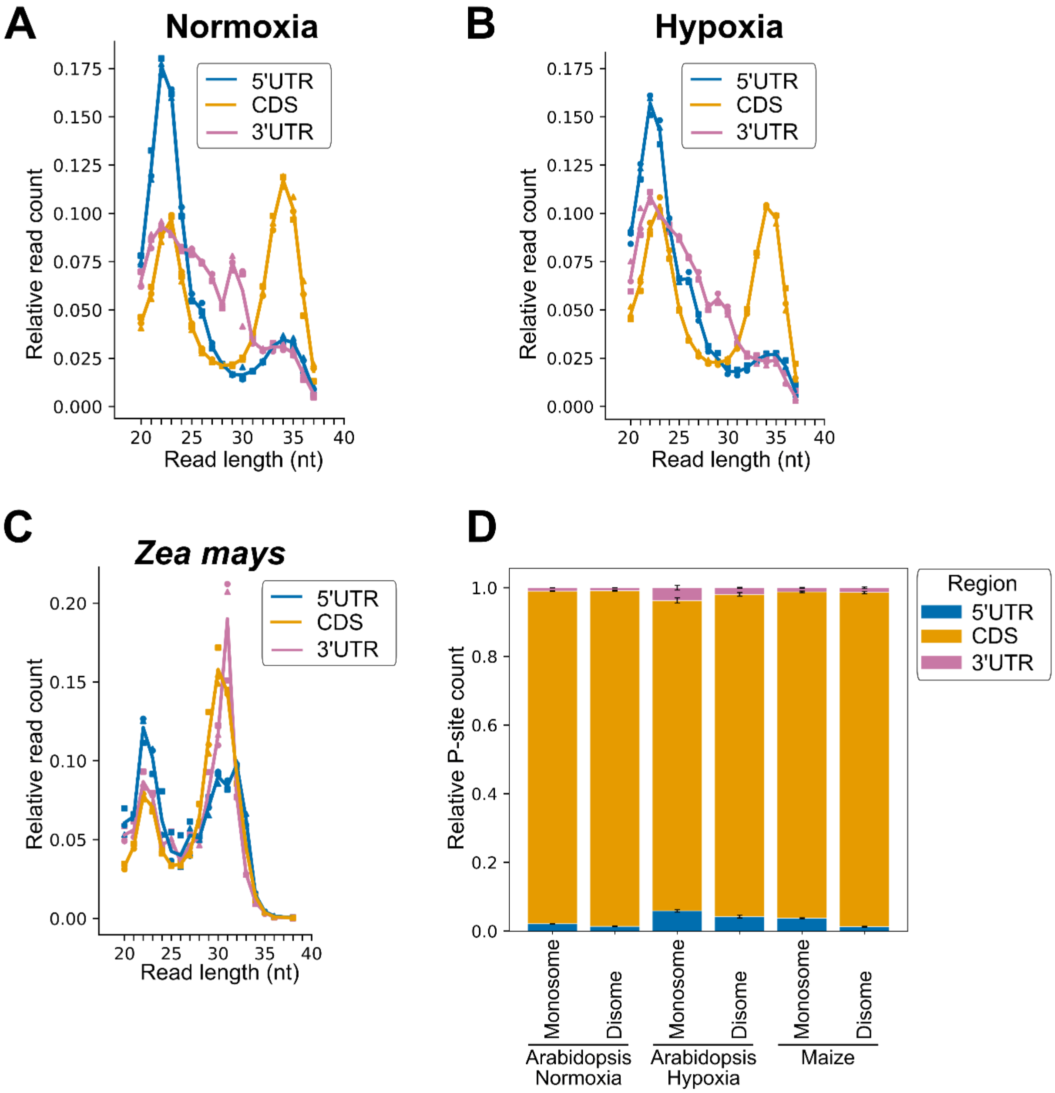


**Fig. S4.** Read length distribution (***A-C***) of monosome-seq footprints on the 5'-UTR, CDS or 3'-UTR identifies high levels of 22-23 nt footprints on the 5'-UTR. *Arabidopsis thaliana* normoxic (***A***) or hypoxic (***B***) seedling monosome-seq data and *Zea mays* shoot data (***C***) are shown. (***D***) Percentage of P-site distribution along the 5'UTR, CDS and 3'UTR.

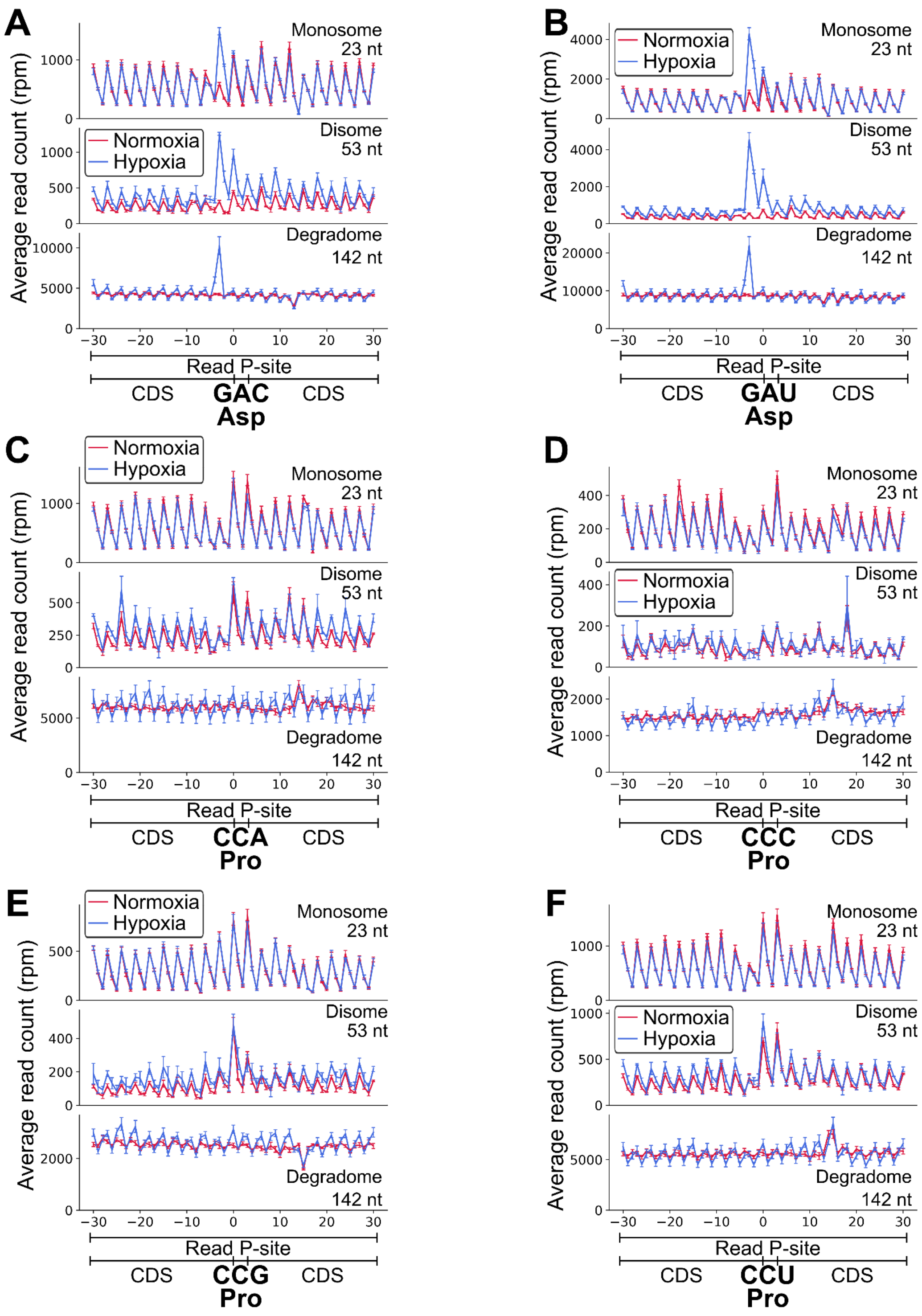


**Fig. S5**. Ribosome P-site RPM distribution around individual Asp and Pro codons. (***A-F***) P-site RPM distribution around Asp GAC (***A***) or GAU (***B***) and Pro CCA (***C***), CCC (***D***), CCG (***E***) or CCU (***F***) codons of different library preparation types and read lengths. Error bars represent standard deviation from three biological replicates.

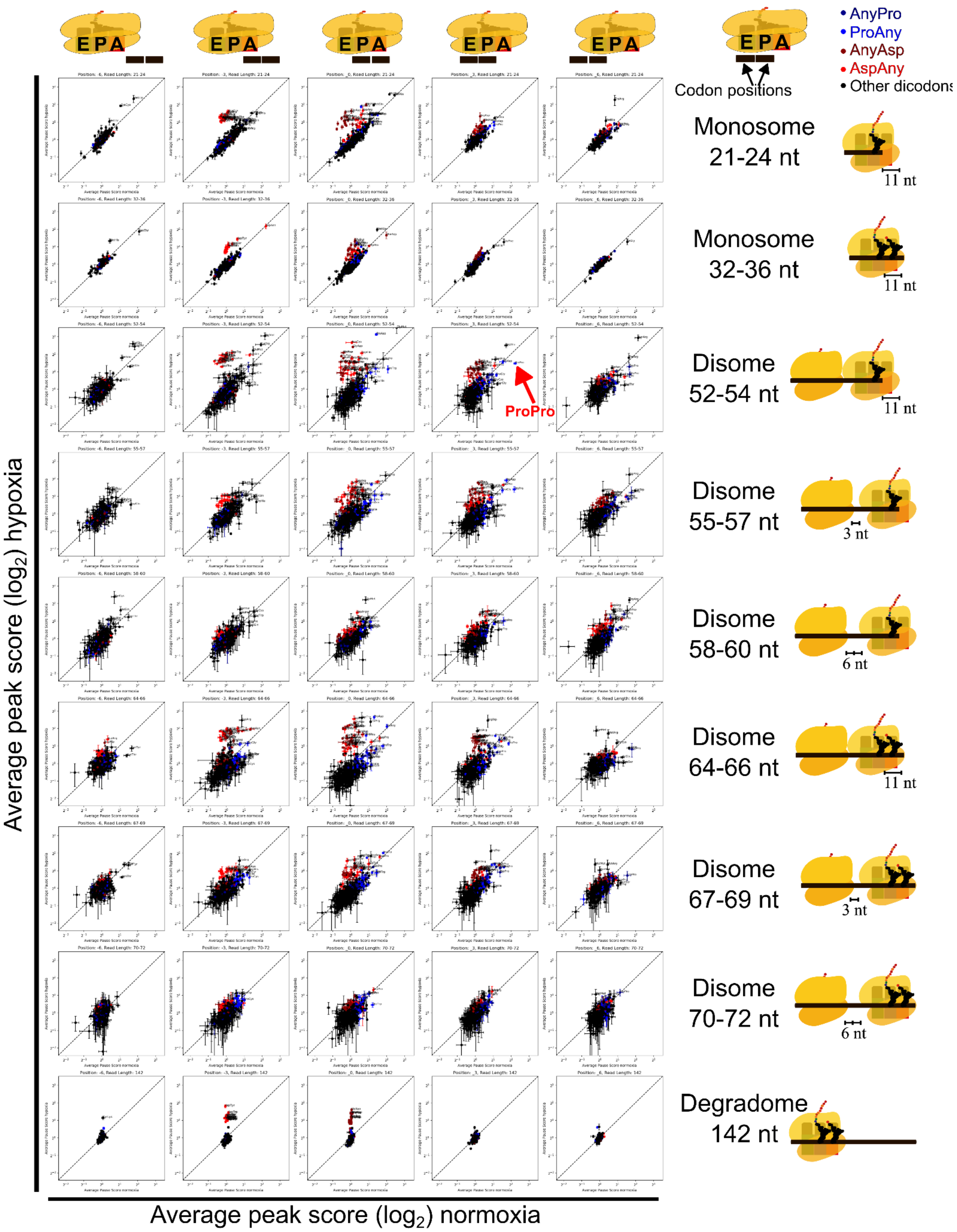


**Fig. S6**. Log₂ peak score profiles of dipeptide codons comparing normoxia (*x*-axis) and hypoxia (*y*-axis) conditions, shown for monosome, disome, and 5'P-degradome-seq datasets. Each column of plots represents the peak scores when dipeptide codons occupy specific positions relative to the ribosome. Each row corresponds to a different ribosomal conformation, as defined by distinct footprint lengths. Data points and error bars represent the mean ± standard deviation from three biological replicates.

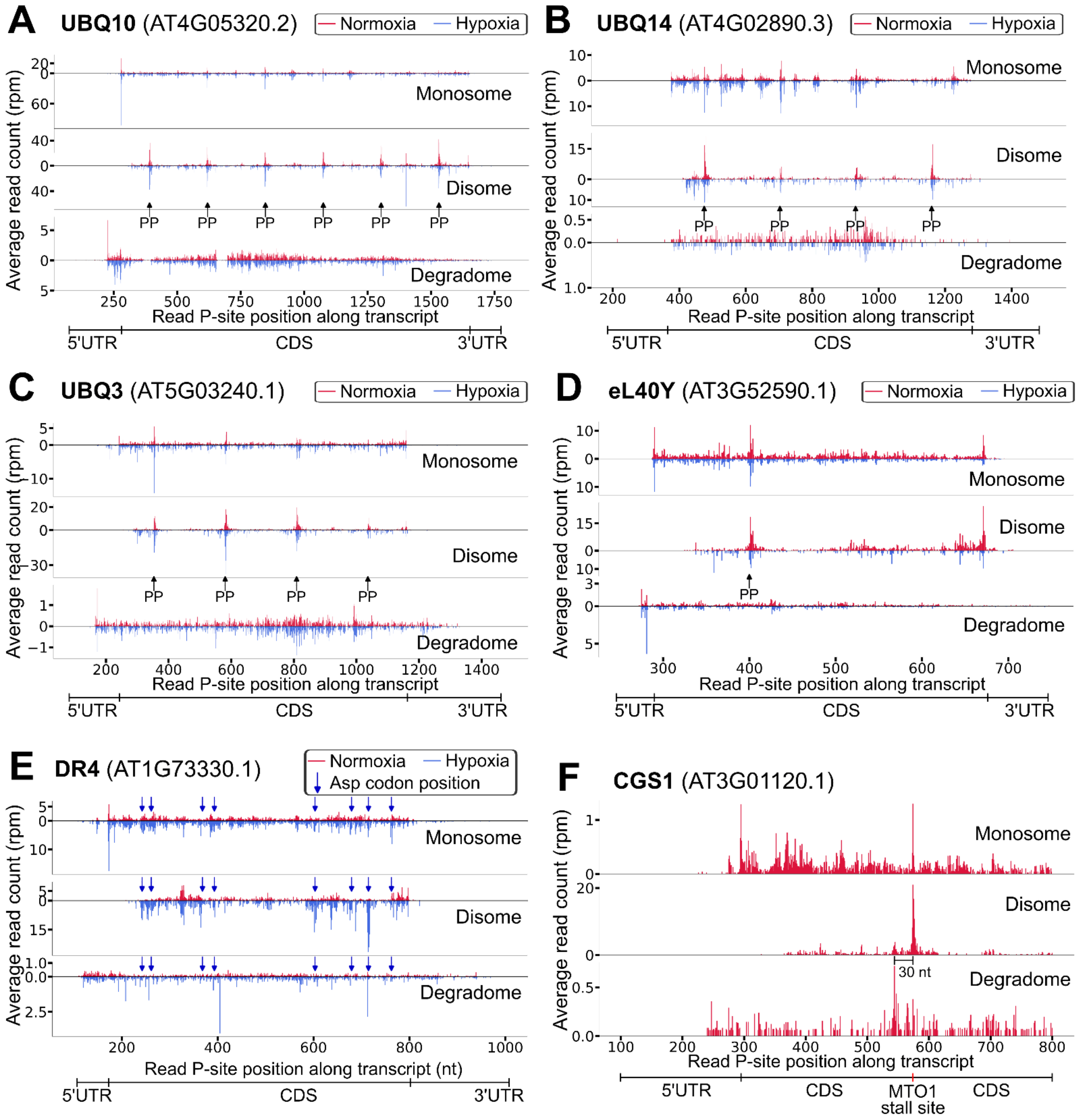


**Fig. S7.** Average read counts (RPM) at predicted ribosomal P-site positions across different transcripts. (***A–D***) Ribosome stalling is observed at di-proline motifs (PP) within ubiquitin domains. Stalling is evident in both monosome and disome data, but not in degradome data. Normoxia data in (***A***) are the same as shown in **Fig. 2E**. (***E***) Example of hypoxia-induced ribosome stalling at aspartate codons. All Asp codons are indicated by blue arrows. *DR4/KT11* encodes a plant-specific protease and is among the 54 translated mRNAs specific to mesophyll cells expressing a key susceptibility gene associated with mesophyll cell transformation into haustoria by the biotroph downy mildew (*Hyaloperonospora arabidopsidis)* (94). (***F***) Ribosome stalling at the known MTO1 site on the *CGS1* transcript. Arabidopsis Gene Identifiers are provided. UBI, *UBIQUITIN;*, eL40Y, *RIBOSOMAL PROTEIN eL40Y;* DR4, *DROUGHT-REPRESSED 4*; CGS1, *CYSTATHIONINE GAMMA-SYNTHASE 1*.

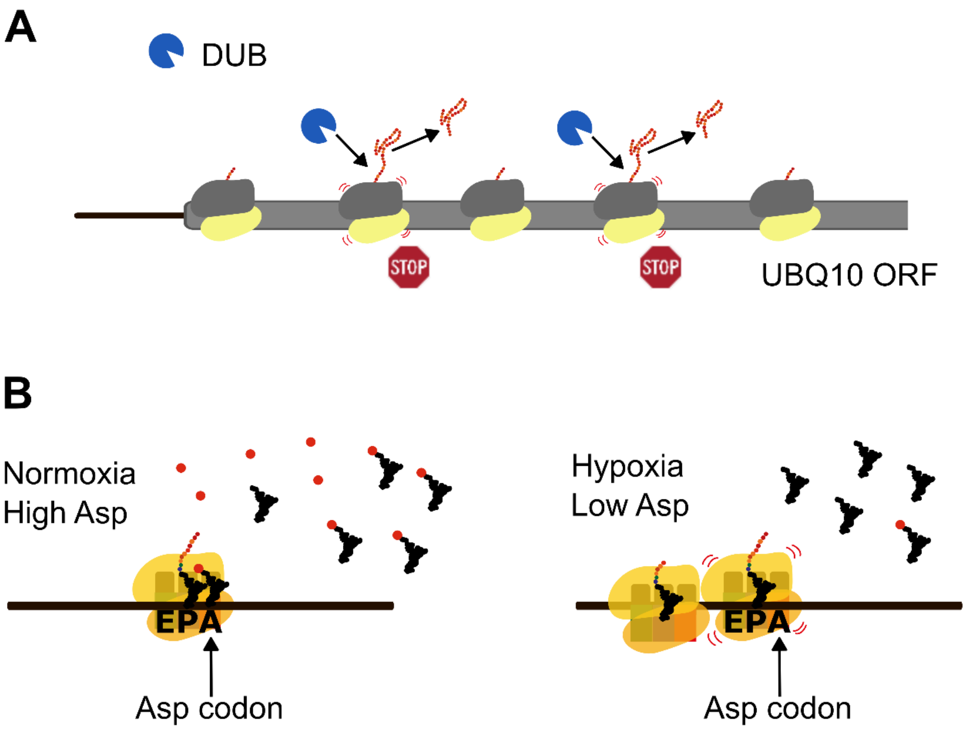


**Fig. S8.** Hypotheses for pronounced ribosome stalling events. (***A***) Di-proline stall sites in polyubiquitin-encoding mRNAs may facilitate co-translational proteolytic cleavage of individual monoubiquitins by de-ubiquitilation enzymes (DUBs). Shown as blue proteases. *A. thaliana* encodes over 60 DUBs but the subgroup involved in processing of ubiquitin monomers from UB Qs has proven challenging due to mutant lethality (95). (***B***) Hypoxia-induced stalling is predominantly observed in ribosomes with a vacant A-site. Given previous reports of reduced aspartate concentrations under hypoxia, we propose the following mechanism: under normoxia, ribosome pausing does not occur due to sufficient levels of charged Asp-tRNAs. Under hypoxia, reduced levels of Asp limit availability of charged Asp-tRNAs, delaying A-site loading, thereby causing ribosome stalling that can lead to ribosome collision-associated co-translational mRNA decay. Whether this mechanism is important for conditional reduction in a transcript or protein is yet to be determined.

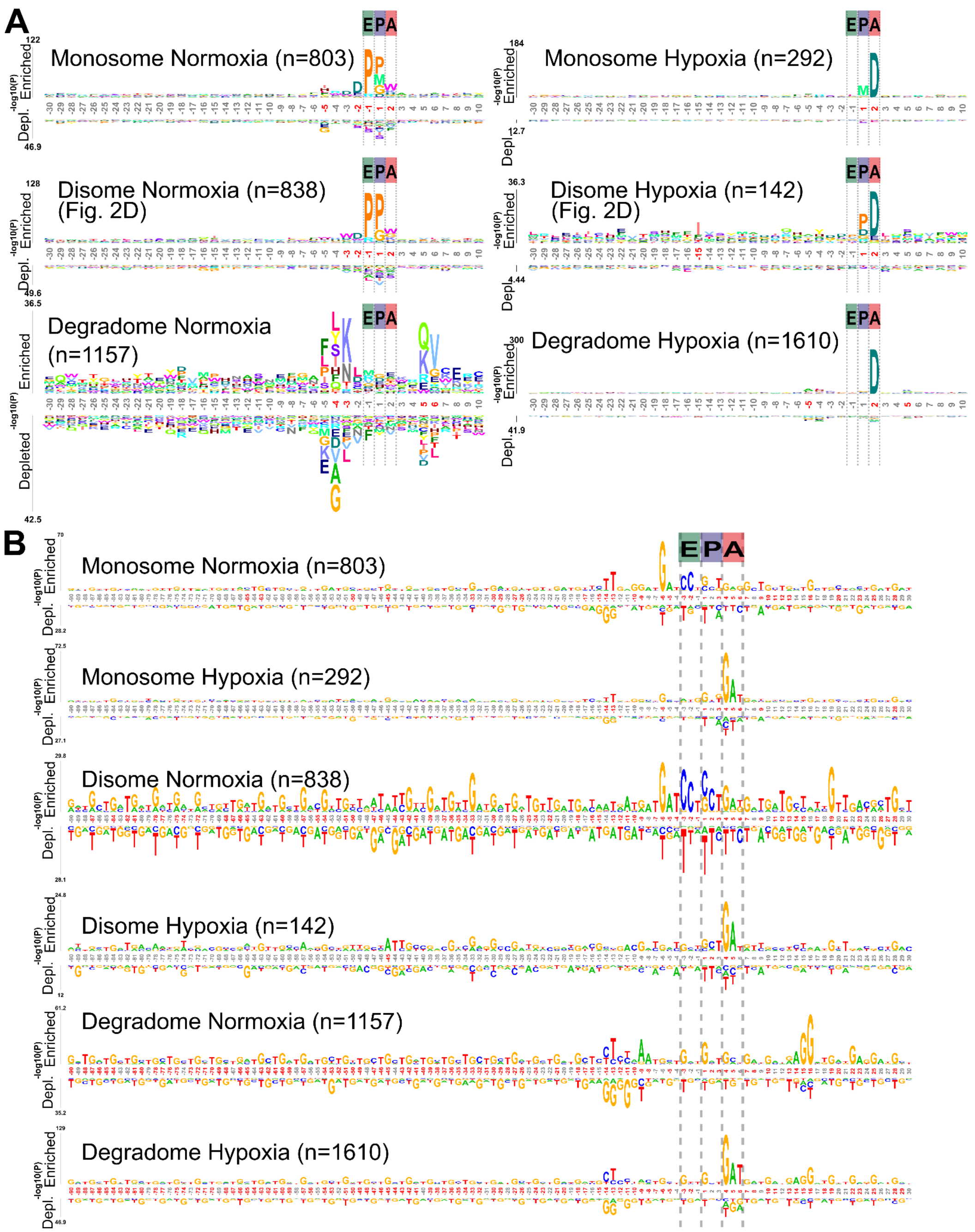


**Fig. S9**. Probability logos of amino acid and mRNA sequences at ribosomal pause sites identified in degradome, monosome, and disome datasets of Arabidopsis. Logos were generated using kpLogo (version 1.0; (86)). Both amino acid-based input (***A***) and nucleotide-based input (***B***) were analyzed. Predicted E-, P-, and A-sites of the stalled ribosomes are indicated. Amino acids or nucleotides shown at the top of each plot indicate enriched residues, whereas those at the bottom indicate depleted residues. The number of input sequences used for each logo is shown in parentheses. Positions highlighted in red represent significantly enriched or depleted motifs (Bonferroni-corrected p-value < 0.01). The bottom two panels of (*A*) correspond to the same data shown in Fig. 2D, but additionally display significantly depleted sequence features. See **File S1** for input sequences.

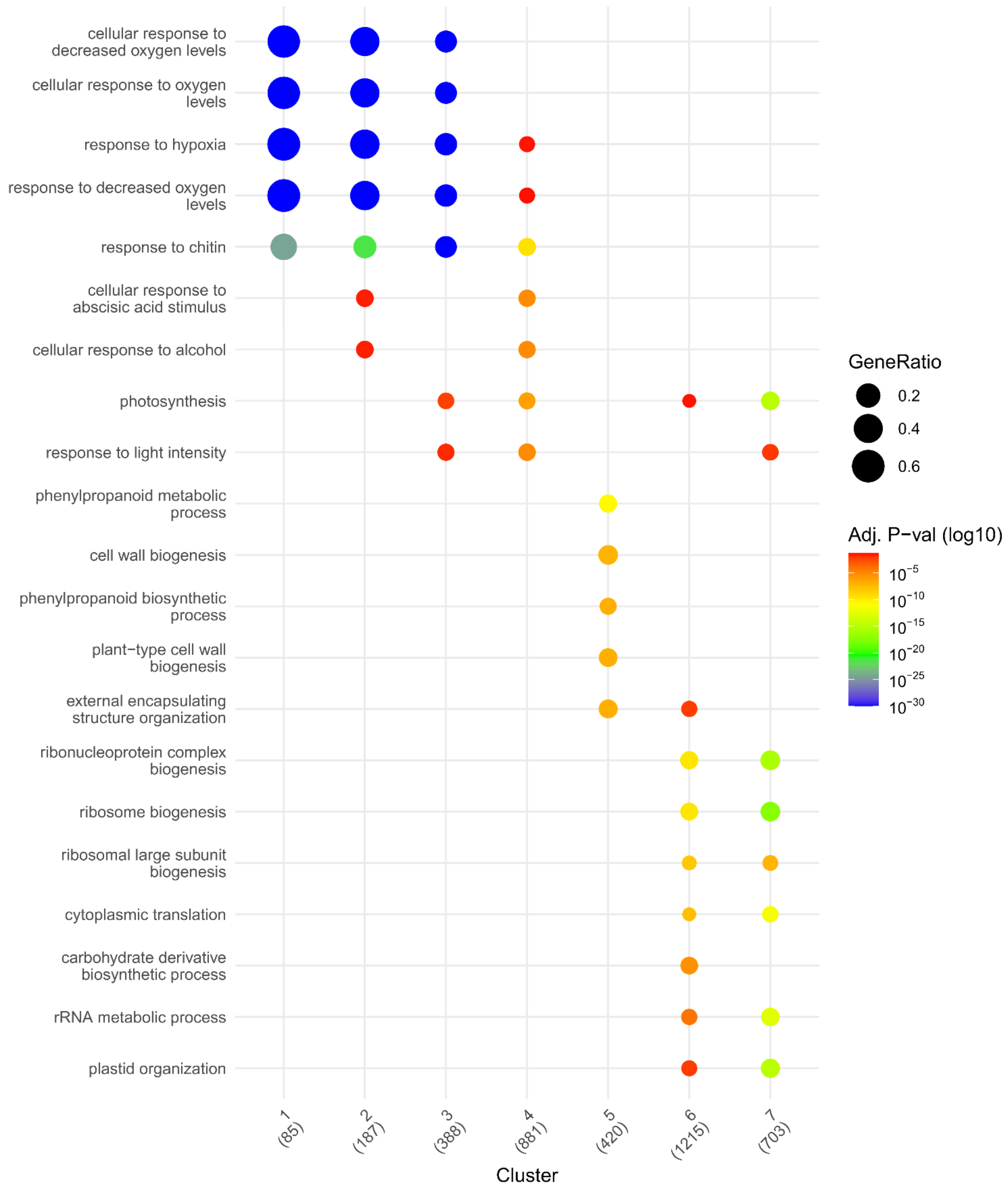


**Fig. S10**. Gene ontology analysis of different clusters in **Fig. 3** using clusterprofiler and Enrichplot R packages for *Arabidopsis thaliana*.

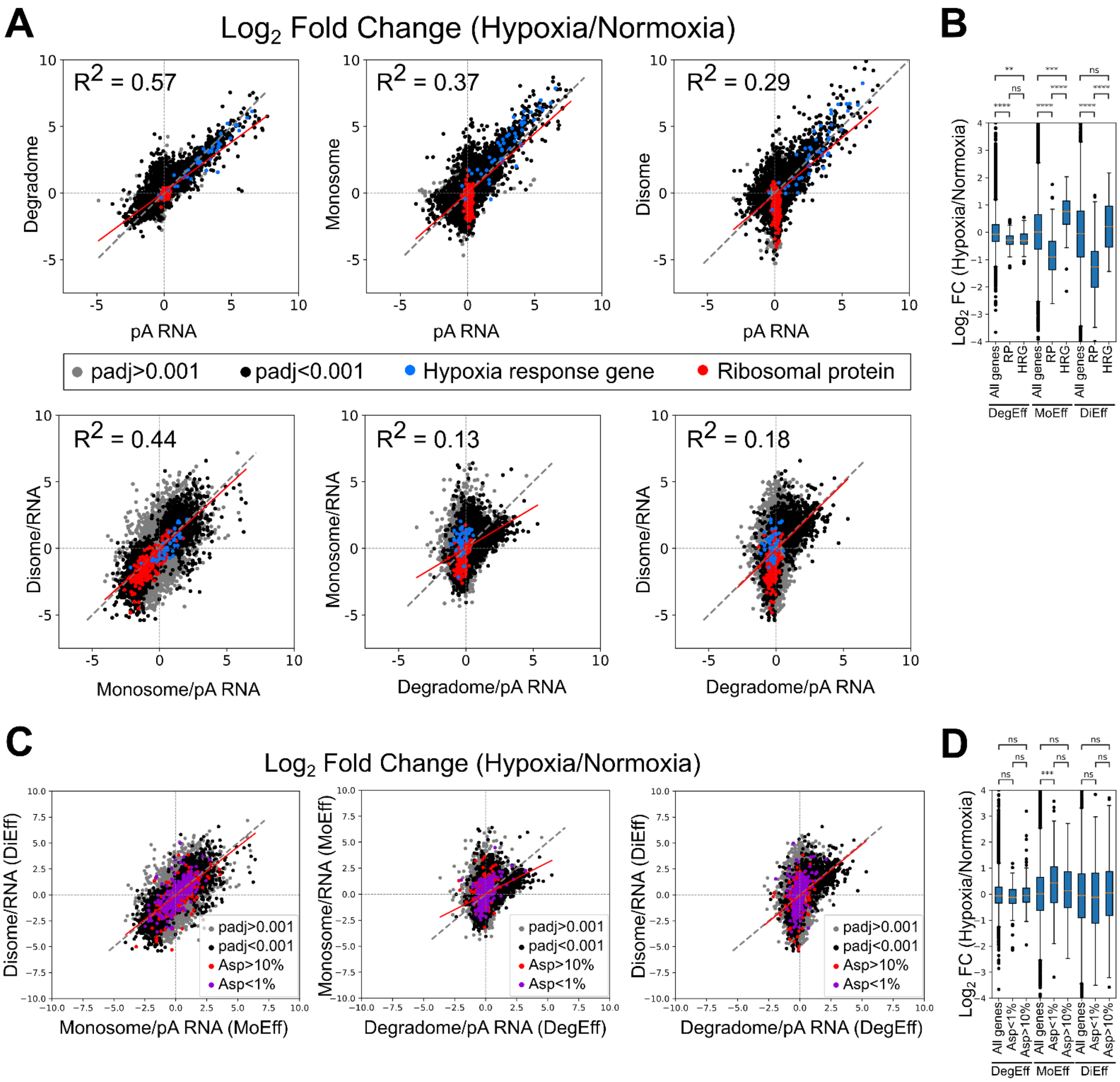


**Fig. S11**. (***A***) Correlation plots of $\text{Log}_2$ FC (Hypoxia/Normoxia) across different data types (top row) or the $\text{Log}_2$ FC of the calculated “translational efficiencies” (normalized against RNA abundance) (bottom row). Hypoxia-response genes and ribosomal proteins are highlighted in blue and red, respectively. Each data point represents the mean of three biological replicates. (***B***) Boxplots summarizing the distributions of $\text{Log}_2$ FC values for the indicated datasets and gene groups. (**C**, **D**) These are the same as (**A, B**) but highlight genes whose coding sequences contain >10% or <1% Asp codons. MoEff (Monosome reads/polyA mRNA reads) (pA) RNA is commonly referred to as Translational Efficiency (TE). DiEff and DegEff are the same calculation but for disome-seq and 5’P-seq reads. Hypoxia Response Genes and Ribosomal Protein genes are identified in **Dataset S2A**.

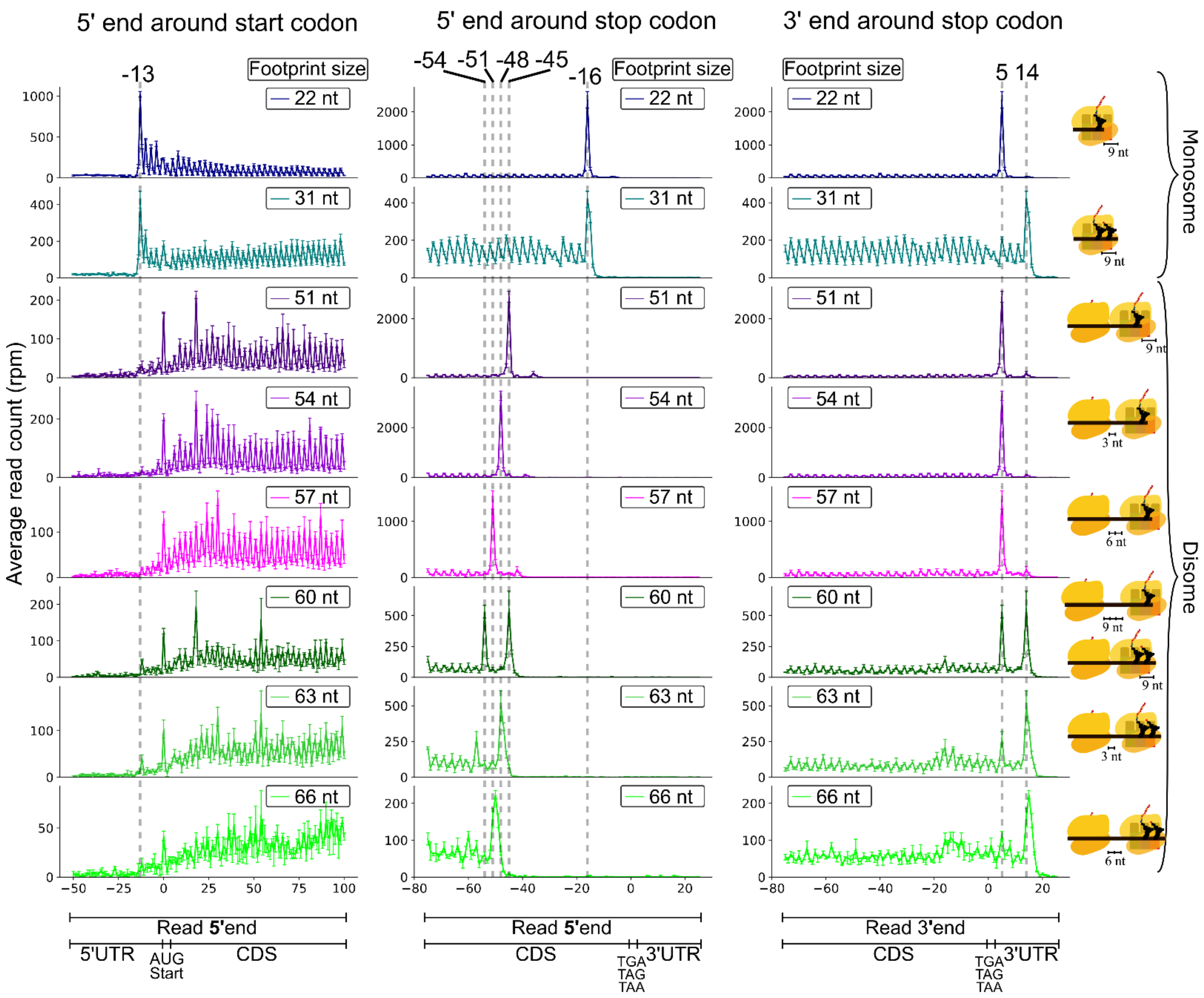


**Fig. S12**. Monosome and disome footprint distribution around the start and stop codon of shoots from vegetative stage V3 B73 maize dataset. Average rpm ± standard deviation from three biological replicates are shown. Illustrations on the far right illustrate the different ribosomal conformations that the different read lengths represent. Note that a read length of 60 nt can represent both 3-codon spaced vacant A-site ribosomes or tightly stacked occupied ribosomes.

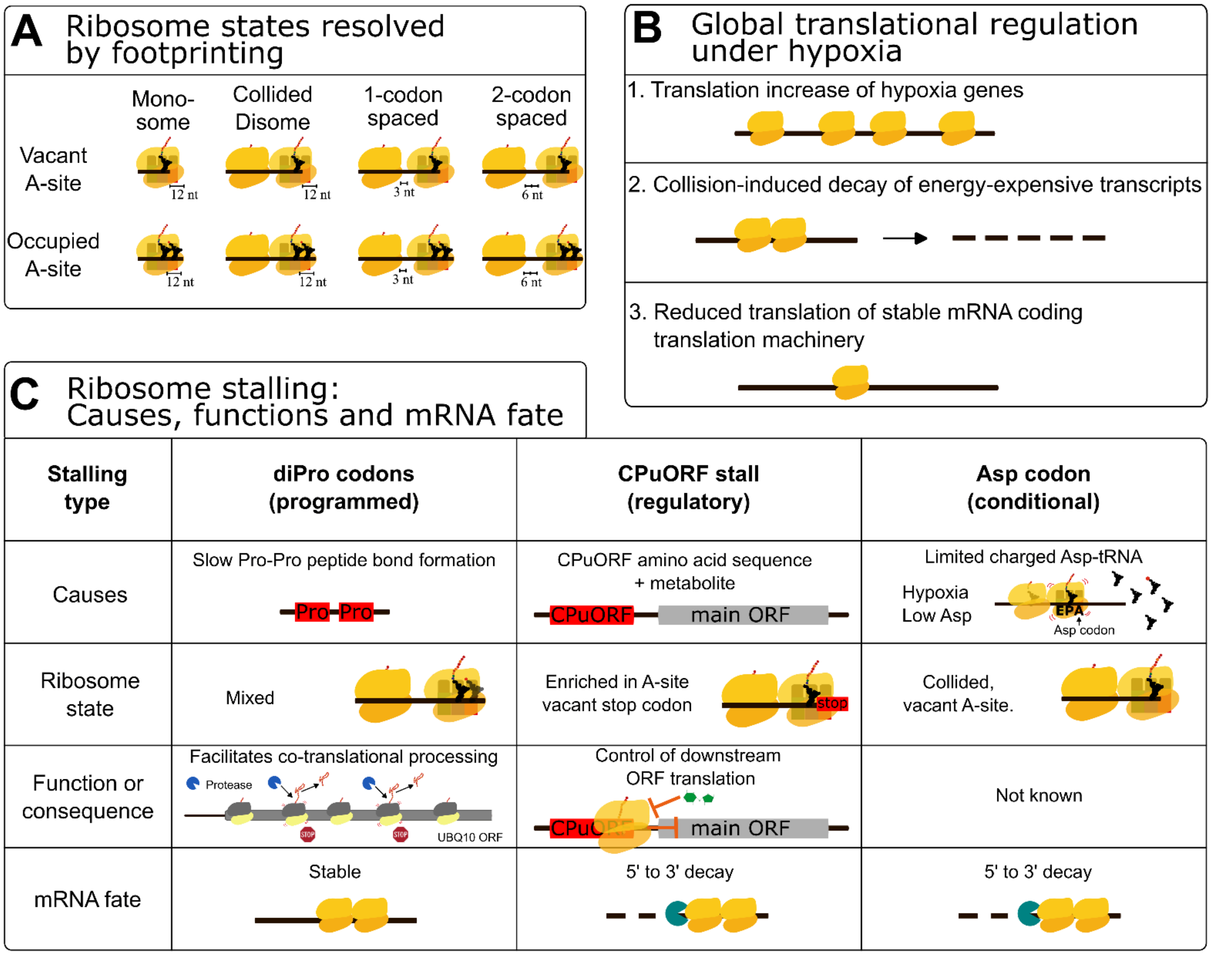


**Fig. S13**. Summary model of the observed molecular causes of ribosome stalling, associated ribosomal states, functional consequences of stalling, and fate of mRNAs undergoing 5' to 3' co-translational decay. (**A**) Combined monosome- and disome-seq enabled mapping of eight distinct ribosomal states at codon resolution along mRNAs. Ribosomes with vacant A-sites produce a small footprint size not typically monitored. (**B**) Global analyses of hypoxia-treated Arabidopsis seedlings, integrating steady-state transcript abundance, translational profiling and free 5′P termini sequencing, revealed multilayered regulation of gene expression: 1. Previously defined Hypoxia Response Genes are translationally upregulated after brief hypoxia. 2. Energy-expensive cell wall-related transcripts exhibit ribosome collisions accompanied by enhancement of co-translational decay. 3. mRNAs encoding components of the translational machinery including most ribosomal protein and translation factors display reduced translation. (**C**) Three stalling types are contrasted: di-Pro codon stalling, CPuORF C-terminal stalling, and Asp codon stalling under hypoxia associated with reduced Asp abundance.

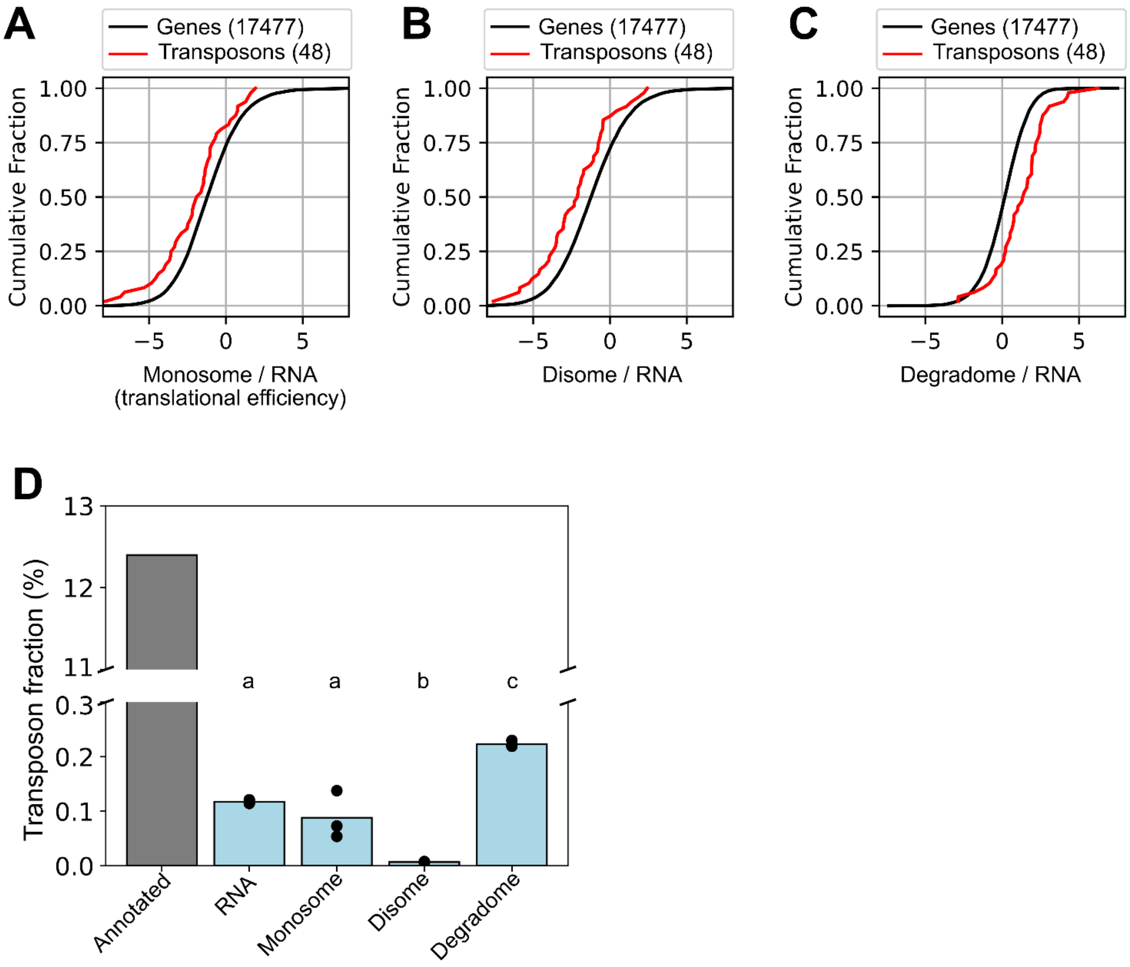


**Fig. S14**. Transposable element transcripts are less efficiently translated and more prone to degradation. (***A-C***) Relative cumulative efficiencies for monosome-seq, disome-seq and degradome-seq datasets of coding sequence (CDS)-containing transcripts (RNA) of annotated transposable elements as compared to all protein-encoding transcripts. Relative efficiencies were calculated by dividing the transcripts per million (TPM) (monosome-seq or disome-seq) or reads per million (RPM) (degradome-seq) by the TPM of RNA-seq. Cumulative fractions of genes or transposable elements that have at least one read in each of the polyadenylated RNA-seq replicates and in at least one of the replicates in any of the monosome, disome and degradome datasets. Numbers in brackets indicate the number of genes or transposable elements used for the analysis. (***D***) Fraction of transposable elements in the data (CDS-containing genes and transposable elements) (annotated) and the fraction of transposable element TPM (RNA-seq, monosome-seq and disome-seq) or RPM (degradome-seq). Statistical analysis performed by one-way ANOVA followed by a Tukey HSD test. Differences of p-value $<0.01$ were considered significant

]

Townsley, Brad T., Michael F. Covington, Yasunori Ichihashi, Kristina Zumstein, and Neelima R. Sinha. 2015. “BrAD-Seq: Breath Adapter Directional Sequencing: A Streamlined, Ultra-Simple and Fast Library Preparation Protocol for Strand Specific mRNA Library Construction.” *Frontiers in Plant Science* 6 (May): 366.

Wilkins, Oscar G., and Jernej Ule. 2021. “Ribocutter: Cas9-Mediated rRNA Depletion from Multiplexed Ribo-Seq Libraries.” In *bioRxiv*. BioRxiv, July 14. https://doi.org/10.1101/2021.07.14.451473.

Willmann, Matthew R., Nathan D. Berkowitz, and Brian D. Gregory. 2014. “Improved Genome-Wide Mapping of Uncapped and Cleaved Transcripts in Eukaryotes--GMUCT 2.0.” *Methods (San Diego, Calif.)* 67 (1): 64–73.

Wu, Hsin-Yen Larry, and Polly Yingshan Hsu. 2022. “A Custom Library Construction Method for Super-Resolution Ribosome Profiling in Arabidopsis.” *Plant Methods* 18 (1): 115.

Wu, Tianzhi, Erqiang Hu, Shuangbin Xu, et al. 2021. “clusterProfiler 4.0: A Universal Enrichment Tool for Interpreting Omics Data.” *Innovation (Cambridge (Mass.))* 2 (3): 100141.

Wu, Xuebing, and David P. Bartel. 2017. “kpLogo: Positional K-Mer Analysis Reveals Hidden Specificity in Biological Sequences.” *Nucleic Acids Research* 45 (W1): W534–W538.

Zhang, Yujie, and Vicent Pelechano. 2021. “High-Throughput 5’P Sequencing Enables the Study of Degradation-Associated Ribosome Stalls.” *Cell Reports Methods* 1 (1): 100001.
